# Genome reconstruction of the non-culturable spinach downy mildew *Peronospora effusa* by metagenome filtering

**DOI:** 10.1101/842658

**Authors:** Joël Klein, Manon Neilen, Marcel van Verk, Bas E. Dutilh, Guido Van den Ackerveken

**Author notes:** these authors contributed equally to the manuscript (Joël Klein, Manon Neilen).

## Abstract

*Peronospora effusa* (previously known as *P. farinosa f. sp. spinaciae*, and here referred to as *Pfs*) is an obligate biotrophic oomycete that causes downy mildew on spinach (*Spinacia oleracea*). To combat this destructive disease resistant cultivars are continually bred. However, new *Pfs* races rapidly break the employed resistance genes. To get insight into the gene repertoire of *Pfs* and identify infection-related genes, the genome of the first reference race, *Pfs1,* was sequenced, assembled, and annotated. Due to the obligate biotrophic nature of this pathogen, material for DNA isolation can only be collected from infected spinach leaves that, however, also contain many other microorganisms. The obtained sequences are, therefore, considered a metagenome. To filter and obtain *Pfs* sequences we utilized the CAT tool to taxonomically annotate ORFs residing on long sequences of a genome pre-assembly. This study is the first to show that CAT filtering performs well on eukaryotic contigs. Based on the taxonomy, determined on multiple ORFs, contaminating long sequences and corresponding reads were removed. Filtered reads were re-assembled to provide a clean and improved *Pfs* genome sequence of 32.40 Mbp consisting of 8,635 scaffolds. Transcript sequencing of a range of infection time points aided the prediction of a total of 13,277 gene models, including 99 RXLR(-like) effector, and 14 putative Crinkler genes. Comparative analysis identified common features in the secretomes of different obligate biotrophic oomycetes, regardless of their phylogenetic distance. Their secretomes are generally smaller, compared to hemibiotrophic and necrotrophic oomycete species. We observe a reduction in proteins involved cell wall degradation, in Nep1-like proteins (NLPs), proteins with PAN/apple domains, and host translocated effectors. The genome of *Pfs1* will be instrumental in studying downy mildew virulence and for understanding the molecular adaptations by which new isolates break spinach resistance.

## Introduction

Phytopathogenic oomycetes are eukaryotic microbes that infect a large range of plant species. Due to their hyphal infection structures they appear fungal-like, however, taxonomically belong to the Stramenopiles [1]. The most devastating phytopathogenic oomycetes are found within the orders of *Albuginales, Peronosporales* and *Pythiales*.

The highly radiated *Peronosporales* order contains species with different lifestyles. The most infamous species of this order are in the hemibiotrophic *Phytophthora* genus. Other species within the *Peronosporales* are the obligate biotrophic downy mildews that cause disease while keeping the plant alive. The relationship between downy mildews and *Phytophthora* species have long been unresolved [2]. In many studies of oomycete phylogeny, the downy mildew species were barely covered. This is mainly because the obligate biotrophic nature of the species makes them hard to work with and they are, therefore, under sampled compared to other oomycete phytopathogens.

The first phylogenetic trees based on morphological traits and single gene comparisons [3, 4] classified the downy mildews as a sister clade to the *Phytophthora* species within the order of Peronosporales. Recently, studies using multiple gene and full genome comparisons suggest that the downy mildews have multiple independent origins within the *Phytophthora* genus [2, 5, 6].

The downy mildew *Peronospora effusa* (previously known as *P. farinosa forma specialis spinaciae*, and here referred to as *Pfs*), is the most important pathogen of spinach. *Pfs* affects the leaves, severely damaging the harvestable parts of the spinach crop. Under favorable environmental conditions, *Pfs* infection can progress rapidly resulting in abundant sporulation within a week post inoculation that is visible as a thick grey ‘furry layer’ of sporangiophores producing abundant asexual spores [7] Preventing spread of this pathogen is difficult, since only a few fungicides are effective in chemical control [8]. As a result, the disease can cause severe losses in this popular crop, and infected fields often completely lose their market value.

During infection, hyphae of *Pfs* grow intercellularly through the tissue and locally breach through cell walls to allow the formation of haustoria [9]. These invaginating feeding structures form a platform for the intimate interaction between plant and pathogen cells, and function as a site for the exchange of nutrients, signals and proteins. Several of the delivered pathogen proteins are known to alter host immunity [10], and subsequently escape plant immune response [11]. These and other molecules secreted by pathogens to promote the establishment and maintenance of a successful infection in the host are called effectors.

Here we describe the sequencing of genomic DNA obtained from *Pfs* spores collected from infected spinach plants using a combination of Illumina and PacBio sequencing. Sequencing of obligate biotrophic species is complicated as the spore washes of infected plant leaves contain many other microorganisms. Bioinformatic filtering on taxonomy using the recently developed Contig Annotation Tool CAT [12] was deployed to remove the majority of contaminating sequences. The obtained assembly of race *Pfs1* was used to predict genes and compare its proteome, in particular its secretome, with that of other oomycete taxa. We show that the secretomes of obligate biotrophic oomycetes are functionally more similar to each other than to that of more closely related species with a different lifestyle.

## Materials and Method

### Downy mildew infection

*Peronospora effusa* race 1 (*Pfs1*) was provided by the Dutch breeding company Rijk Zwaan Breeding BV in 2014. As *Pfs1* is an obligate biotrophic maintenance was done on *Spinacia oleracea* Viroflay plants. Seeds were sown on soil, stratified for two days at 4**°**C and grown under long day condition for two weeks (16h light, 70% humidity, 21°C). sporangiophores were washed off infected plant material in 50 ml falcon tubes. The solution was filtered through miracloth and the spore concentration was checked under the microscope. Four-day-old *Spinacia oleracea* Viroflay plants were infected with *Pfs* by spraying a spore solution (70 spores/ul) in tap water. Seven days post inoculation, *Pfs* sporangiospores were collected from heavily-infected spinach leaves with tap water, using a soft brush to prevent plant and soil contamination and used for DNA isolation and genome sequencing.

### DNA isolation and genome sequencing

The sporangiospores were freeze dried, ground and dissolved in CTAB (Cetyltrimethyl ammonium bromide) extraction buffer, lysed for 30 minutes at 65 **°**C, followed by a phenol-chloroform/isoamyl-alcohol, and chloroform/isoamyl-alcohol extraction. DNA was precipitated from the aqueous phase with NaOAc and ice-cold isopropanol. The precipitate was collected by centrifugation, and the resulting pellet washed with ice cold 70% ethanol. DNA was further purified using a QIAGEN Genomic-tip 20/G, following the standard protocol provided by the manufacturer. DNA was quantified using a Qubit HS dsDNA assay (Thermo Fisher Scientific) and sheared using the Covaris S220 ultrasonicator set to 550 bp. The sequencing library was constructed with the Illumina TruSeq DNA PCR-Free kit. Fragment size distribution in the library was determined before and after the library preparation using the Agilent Bioanalyzer 2100 with HS-DNA chip (Agilent Technologies). The library was sequenced on an Illumina Nextseq machine in high output mode with a 550 bp genomic insert paired end 150 bp reads. Illumina reads with low quality ends were trimmed (Q<36) using prinseq-lite [13].

For PacBio sequencing the input DNA was amplified by WGA (Whole Genome Amplification) using the Illustra GenomiPhi V2 DNA Amplification (GE Healthcare). The sequencing library for PacBio was constructed according to the manufacturer protocol. The resulting library was sequenced on 24 SMRT cells (P6 polymerase and C4 chemistry) using the RSII sequencer. The obtained PacBio reads were error-corrected using the Falcon pipeline [14] with the standard settings using the SMRT Portal that is part of the SMRT analysis software package version 2.3.0 from PacBio [15]. The analysis software package was installed according to the installation instructions on an Amazon WebService (AWS) cloud-based computer and operated via its build in GUI.

### Taxonomic classification of long reads

The taxonomic origin of each error corrected PacBio read was determined using the CAT (Contig Annotation Tool) pipeline version 1.0 with default parameters [12]. To do this, CAT first identifies open reading frames (ORFs) on the long sequences or contigs using Prodigal [16] and queries them against the NCBI non-redundant (nr) protein database (retrieved November 2016) using DIAMOND [17]. A benchmarked weighting scheme is then applied that allows the contig to be classified with high precision [12].

### Genome assembly and identification of repeats

A pre-assembly was made using taxonomically filtered and corrected PacBio sequences and 60% of the Illumina reads using SPAdes version 3.5.0 [18]. The error-corrected PacBio reads were used as long reads in the assembly, SPAdes was set to use *k*-mer lengths of: 21, 33, 55, 77, 99, 127 for the assembly and the --careful option was used to minimize the number of mismatches in the final contigs. The contigs derived from the pre-assembly were filtered using the CAT tool (see above), and sequences that were designated as bacterial or non-stramenopile eukaryotes were collected. The entire set of Illumina sequencing reads were aligned to the collection of removed sequences (annotated as bacterial and non-stramenopile) with Bowtie version 2.2.7 using default settings [19]. Illumina reads that aligned to these sequences were removed from the Illumina data set. The remaining Illumina reads (Illumina filtered), and PacBio sequences were re-assembled with SPAdes (same settings as the preassembly), which resulted in a final *Pfs1* genome assembly. A custom repeat library for the *Pfs1* genome assembly was generated with RepeatModeler [20]. Repeat regions in the assembled *Pfs1* genome were predicted using RepeatMasker 4.0.7 [20].

### Quality evaluation of the assembly

*k-m*ers of length 21 in the filtered Illumina data set were counted with Jellyfish count version 2.0 [21] with settings -C -m 21 -s 1000000000 followed by Jellyfish histo. The histogram was plotted with GenomeScope [22] to produce a graphical output and an estimate of the genome size. The coverage of the genome by PacBio sequences was determined by aligning the unfiltered error-corrected PacBio reads to the *Pfs1* genome assembly using BWA-mem [23] and selected –x pacbio option. The BBmap pileup [24] script was used to determine the percentage covered bases by PacBio reads in the final assembly of the *Pfs1*.

The GC-content per contig larger than 1kb was calculated using a Perl script [25]. GC density plots were generated in Rstudio version 1.0.143 using GGplot version 3.1 [26]. For comparison, the same analysis was done on a selection of other publicly available oomycete assemblies; *Hyaloperonospora arabidopsidis* [27], *Peronospora belbahrii* [28], *Phytophthora infestans* [29], *Bremia lactucae* [30], *Phytophthora parasitica* [31], *Phytophthora ramorum* (Pr102) [31], *Phytophthora sojae* [32], *Peronospora tabacina* (968-S26) [33] and *Plasmopara viticola* [5].

Kaiju [34] was used to analyze the taxonomic origin by mapping reads to the NCBI nr nucleotide database (November 2017). The input for Kaiju was generated using ART [35] set at 20x coverage with 150 bp Illumina to create artificial sequencing reads from the various FASTA assembly files of the genomes of different oomycetes.

Genome completeness and gene duplications were analyzed with BUSCO version 3 [36] with default settings using the protists Ensembl database (May 2018).

### RNA sequencing and gene model prediction

RNA of *Pfs1* at different stages during the infection was isolated and sequenced to aid gene model prediction. Infected leaves and cotyledons were harvested every day from three days post infection (dpi) until sporulation (7 dpi). Besides these infected leaves, spores were harvested, and a subset of these spores were placed in a petri dish with water and incubated overnight at 16 ° C to allow them to germinate. RNA was isolated using the RNeasy Plant Mini Kit from Qiagen, and the RNA was analyzed using the Agilent 2100 bioanalyzer to determine the RNA quality and integrity. The RNA-sequencing libraries were made with the Illumina TruSeq Stranded mRNA LT kit. Paired-end 150 bp reads were obtained from the different samples with the Illumina Nextseq 500 machine on high output mode. RNA-seq reads from all the samples were pooled, aligned to the *Pfs1* assembly using Tophat [37], and used as input for gene model prediction using BRAKER1 [38]. The obtained gene models for the *Pfs1* genome together with the RNA-seq alignment result, the repeat models, and results obtained from a BLAST-p search to the nr NCBI database (January 2017), were loaded into a custom made WebApollo [39] instance. Gene models on the 100 largest contigs of the genome were manually curated and all gene models were exported from the WebApollo instance for further use.

### Gene annotation and the identification of functional domains

Bedtools intersect version 2.27 was used to determine the overlap between *Pfs1* gene models and annotated repeat elements in the genome. Gene models that had more than 20% overlap with a region marked as a repeat-containing gene. ANNIE [40] was used to annotate proteins on the *Pfs1* genome based on Pfam domains [41] and homologous sequences in the NCBI-Swissprot database (accessed Augustus 2017). Sequences that were annotated as transposon*s* by ANNIE were removed from the gene set. SignalP 4.1 [42] was used to predict the presence and location of a signal peptide, the D-cutoff for noTM and TM networks were set at 0.34 to increase sensitivity [43]. TMHMM version 2 [44] was used to predict the presence of transmembrane helices in the proteins of *Pfs1*. To identify proteins that possess one or more WY domains an HMM model made by Win *et al*. [45] was used. Protein sequences that possessed a WY domain were extracted and realigned. This alignment was used to construct a new HMM model using HMMER version 3.2.1 [46] and queried again against all protein models in the *Pfs* genome to obtain the full set of WY domains containing proteins.

### Effector identification

Putative effectors residing on the genome of *Pfs1* were identified with a custom made pipeline [47] constructed using the Perl [48] scripting language. Secreted proteins were screened for the occurrence of known translocation domains within the first 100 amino acids after the signal peptide. Proteins with a canonical RxLR, or a degenerative RxLR (xxLR or RxLx) combined with either an EER-like or a WY domain or both where considered putative RxLR effectors. A degenerative EER domain was allowed to vary from the canonical EER by at most one position.

Proteins with a canonical LFLAK motif or a degenerative LFLAK and HVL motif in the first 100 amino acids of the protein sequence. A HMMer profile was constructed based on the LFLAK or HVL containing proteins. This HMMer profile was used to identify Crinkler effector candidates lacking the LFLAK or HVL motif.

Proteins with an additional transmembrane domain or a C-terminal ER retention signal (H/KDEL) were removed. WY domains were identified using hmmsearch version 3.1b2 [49] with the published *Phytophthora* HMM model (see above) [50]. *Pfs* WY-motif containing protein sequences were realigned and used to construct a *Pfs* specific WY HMM model using hmmbuild version 3.1b2 [49]. Based on the *Pfs* specific HMM model WY-motif containing *Pfs* proteins were determined.

The effector prediction for the comparative analysis was done in a similar fashion, except the published *Phytophthora* HMM model for RxLR prediction and a published model for CRN prediction was used [51]. The prediction of effectors using the same model in each species enabled the comparison.

### Comparative gene distance analysis

Based on the gene locations encoded in the GFF file the 3’ and 5’ intergenic distances between genes on contigs were calculated as a measure of local gene density. When a gene is located next to beginning or end of a contig, the distance was taken from the start or end of the gene to the end of the contig. Putative high confidence RxLR effector sequences that encode for proteins with either an exact canonical RxLR motif or an RxLR-like motif in combination with one or more WY-motifs were selected for the comparison (66 in total). Distances were visualized using a heat map constructed with the GGPlot geom_hex function [26]. Statistical significance was determined using the Wilcoxon signed-rank test [52].

### Comparative secretomics

The predicted proteomes of eighteen plant pathogenic oomycetes were obtained from Ensembl and NCBI (S1 Table). Proteins in the collected proteomes that have a predicted secretion signal [42] (SignalP v.4.1, D-cutoff for SignalP-noTM and TM networks = 0.34 [43]), no additional transmembrane domain (TMHMM 2.0 [44]) or C-terminal K/HDEL domain were considered secreted. Functional annotations of the secreted proteins were predicted using InterProScan [53] and the CAZymes database [54] using the dbCAN2 meta server [55].

### Phylogenetic analysis

The phylogenetic relationships between the proteomes of the studied species were inferred using Orthofinder [56]. Orthofinder first identifies ‘orthogroups’ of proteins that descended from a single ancestral protein. Next it determines pairwise orthologs between each pair of species. Orthogroups with only one protein of each species were used to make gene trees using MAFFT [57]. The species tree was inferred from the gene trees using the distance algorithms of FastMe [58] and visualized using EvolView v2 [59].

### Principal Component Analysis

The total number of InterPro and CAZymes domain per species was summarized in a counts table. For each domain the number was divided by the total number of domains for that species. The normalized matrix has been loaded into Phyloseq version 1.22.3 [60] with R version 3.4.4 [61] in RStudio [62]. A PCA plot has been made with the Phyloseq ordinate function on euclidean distance. The PCA plot has been made with the GGPlot R package [26]. The bi plot has been generated with the standard prcomp function in R with the same normalized matrix. Figures were optimized using Adobe Photoshop 2017.01.1.

### Permutational analysis of variance (PERMANOVA)

A PERMANOVA using distance matrices was used to statistically test whether there is a difference between the clades based on their CAZymes and InterPro domains. PERMANOVA is a non-parametric method for multivariate analysis of variance using permutations. The data has been double root transformed with the vegdist function from the R-package vegan version 2.5-3 [63]. After the transformation the PERMANOVA has been calculated with the adonis function from the Vegan package. A total number of 999 permutations have been made to retrieve a representative permutation result.

### Enrichment analysis

A chi-square test with Bonferroni correction was used to identify under- and over-represented InterPro domains in each group (*Hyaloperonospora/Peronospora, Plasmopara, Albugo*) compared to *Phytophthora*. The actual range was the sum of the proteins that have a given domain. The expected range was the fraction of proteins with a given domain that is expected to belong to a species cluster giving the overall ratio of InterPro domains between species clusters.

## Results

An early isolate of *Peronospora effusa*, race 1 (*Pfs1)*, was used to create a reference genome as it predates resistance breeding in spinach and its infection is effectively stopped by all spinach resistance genes known to date. Since downy mildews cannot be grown axenically we isolated asexual sporangiospores by carefully washing highly-infected leaves of the universally susceptible cultivar Viroflay. Genomic DNA was isolated from freeze-dried spores and used to construct libraries for PacBio and Illumina sequencing, resulting in 1.09 million PacBio reads with a N50 of 9,253 bp, and 535 million Illumina reads of 150 bp. The paired-end Illumina reads were used for a trial assembly using Velvet. Inspection of the draft assembly showed that many contigs were of bacterial instead of oomycete origin. This is likely caused by contamination of the isolated *Pfs* spores with other microorganisms that reside on infected leaves and that are collected in the wash-offs. We, therefore, decided to treat the sequences as a metagenome and bioinformatically filter the sequences and corresponding reads.

### Taxonomic filtering

To filter out the sequences that could be classified as contaminants we deployed CAT [12] on long reads and contigs derived from assemblies (outlined in the flow diagram in Fig 1). Details on the CAT method are described in the materials and methods section. In short, CAT utilizes the combined taxonomic annotations of multiple individual ORFs found on each sequence to determine its likely taxonomic origin. This allows for a robust taxon classification that is based on multiple hits, rather than a single best hit. An example of the CAT taxonomic classification for two of our sequences is visualized in Fig 2.

**Fig 1.**
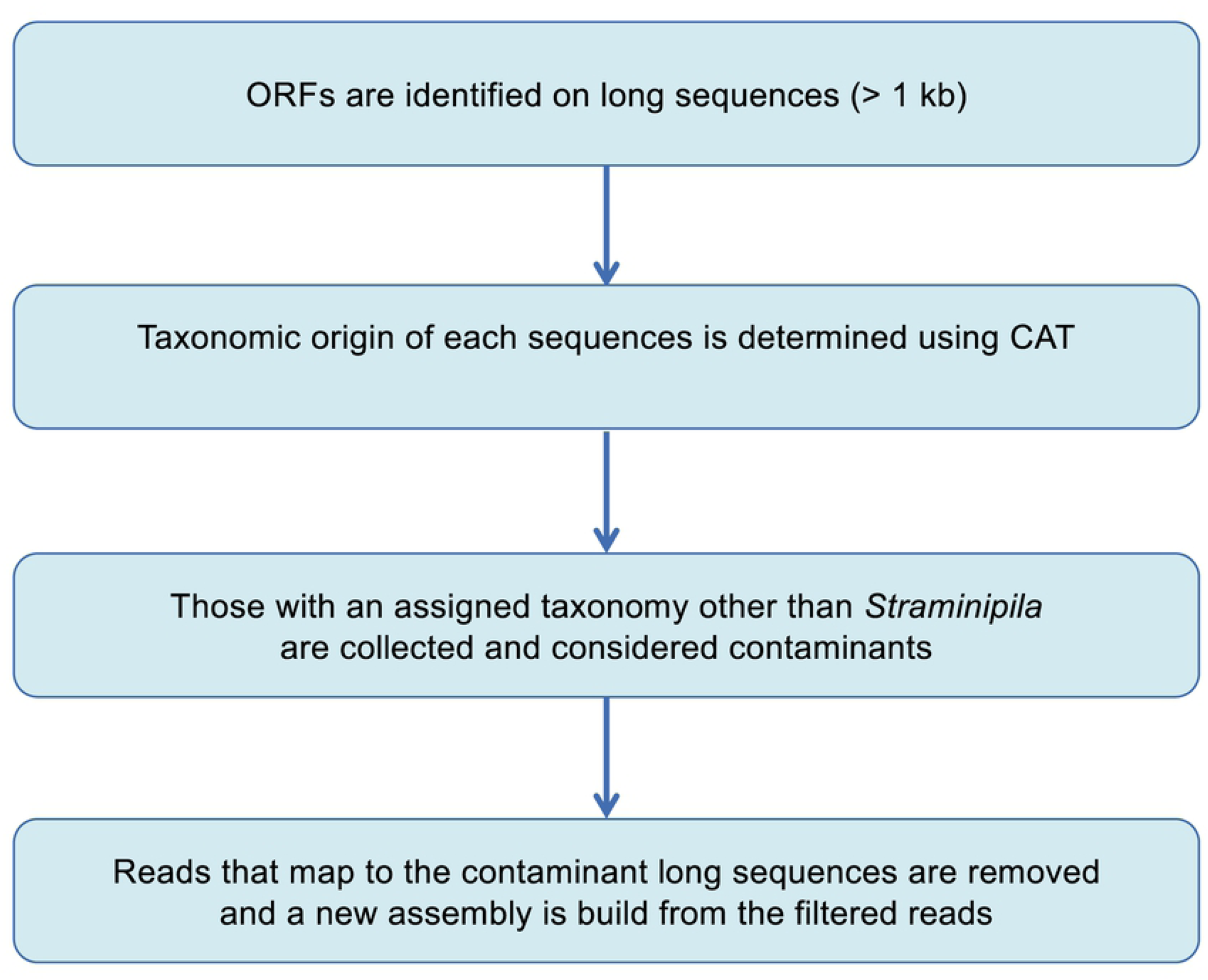
Flow diagram depicting the steps in the taxonomic filtering of long sequences (PacBio reads or assembled contigs).

**Fig 2.**
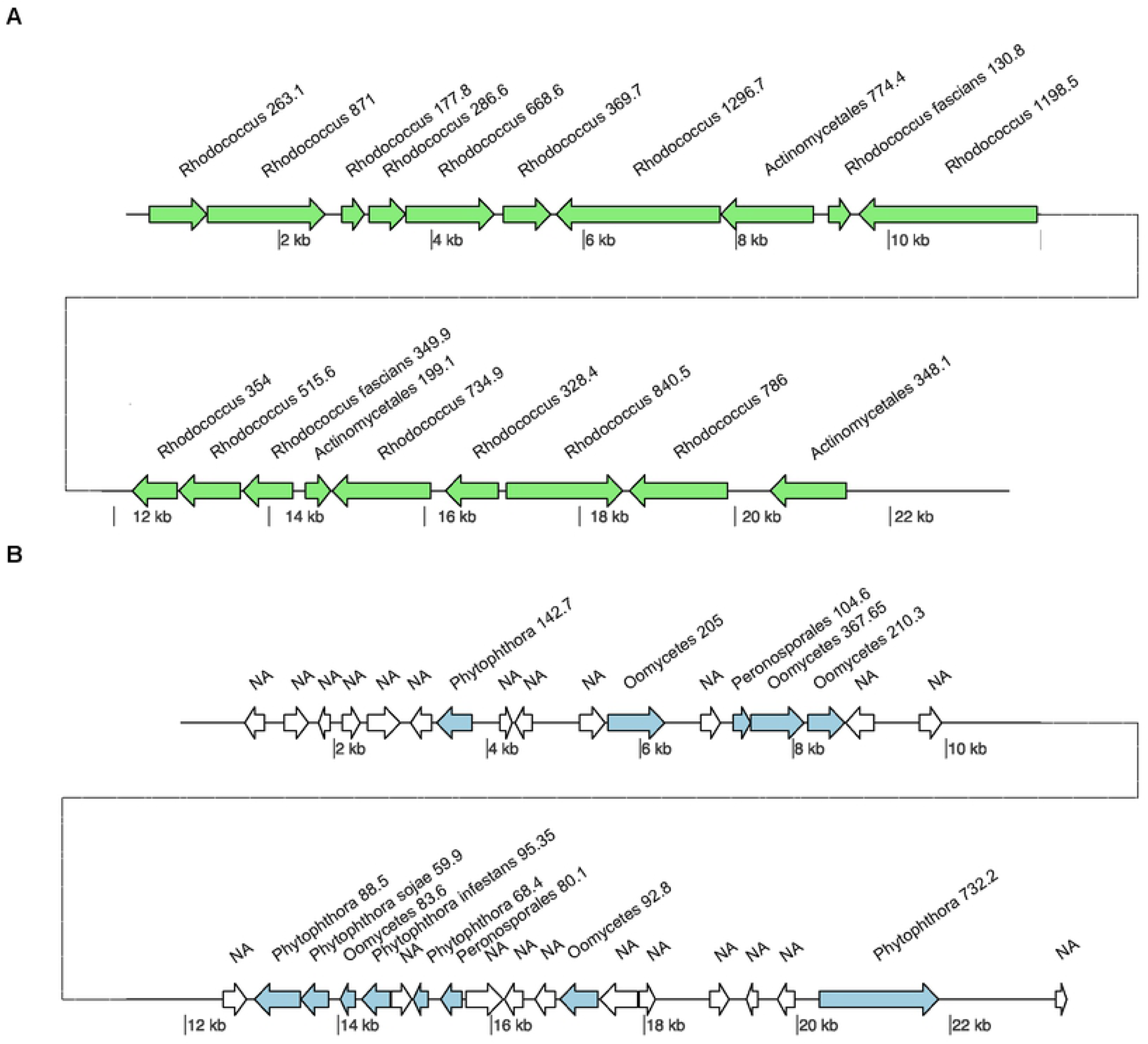
Taxonomic classification by CAT. Two contigs are depicted and per ORF a single top hit is shown. **A** Contig from the pre-assembly assigned by the CAT tool as bacterial, ORFs of bacterial origin are colored green, and ORF with no hits to the database are colored white. On this contig most ORFs had a highest blast hit with *Rhodococcus* species. The ΣBmax for this contig is 10982. and the highest ΣBtaxon is for the *Rhodococcus* genus at 9660, which is well above the cutoff of 5491 (ΣBmax * 0.5). The taxonomic origin of this contig was therefore assigned to the genus *Rhodococcus*, and as a consequence this contig was regarded as non-*Pfs* and removed. **B** Contig from the pre-assembly assigned by the CAT tool as an oomycete contig. On this contig all ORFs have a best hit to an oomycete species, and the ΣBmax is 2328. In fact, most ORFs have a best hit to species in the *Phytophthora* genus (ΣBtaxon :1184), or the Peronosporales family (ΣBtaxon :184). The ΣBtaxon for the *Phytophthora* genus is above the cutoff at 1164 (ΣBmax *0.5) thus assigning this contig to the *Phytophthora* genus, and consequently this contig is maintained for the *Pfs* genome assembly.

CAT was first used on the long PacBio reads. As these reads contain about 15% base call errors on average, they were first error-corrected using the FALCON pipeline. The FALCON pipeline fixes long PacBio reads by mapping short reads obtained in the same runs. The resulting 466,225 PacBio reads had a total length of 1,003 Mb with a N50 of 3,325 bp and were subsequently assigned a taxonomic classification using CAT. PacBio reads that were classified as prokaryotic, or non-stramenopile eukaryotic (e.g. Fungi) were removed, whereas reads with the assigned taxonomy “stramenopiles” or “unknown” were retained.

This resulted in a cleaned set of 232,846 PacBio reads with a total length of 522 Mb with a N50 of 3,458 bp that was used for a hybrid pre-assembly. In order to evaluate the effectiveness of the CAT tool in removing contaminating genomic sequences we analyzed the GC-content of the reads. The corrected PacBio reads showed two distinct peaks (Fig 3a), whereas oomycete genomes have a rather GC band-width around 50%, as shown in S1a Fig for the contigs of the *Phytophthora infestans* genome [29]. After CAT filtering a single peak remained with a narrow GC-content distribution around ∼48%, demonstrating that the tool, that does not take into account GC-content but uses a weighting scheme based on protein sequence similarity, was effective in removing contaminating sequences (Fig 3b).

**Fig 3.**
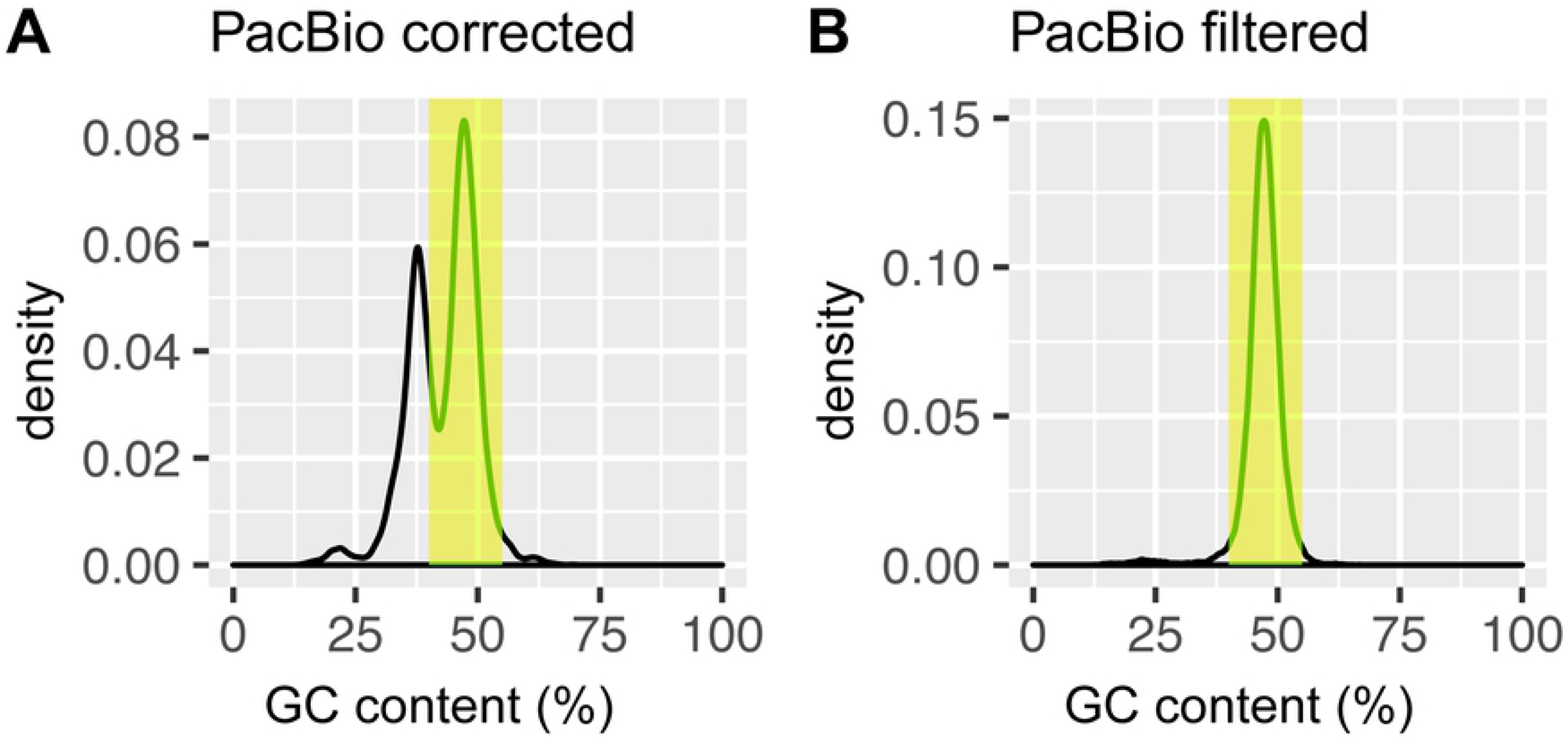
Density plot of the GC values of assembled contigs from the *Pfs1* assembly, and of the PacBio reads before and after CAT filtering sequences. The green bar indicates the region between 40 and 55% GC, based on reads >1 kb. **A.** PacBio reads before CAT-filtering show a bimodal distribution with a presumed peak of contaminating sequences with a GC content of ∼40 %. **B.** PacBio reads after CAT-filtering show a distribution consisting of a single peak with a GC content around ∼46%.

### Hybrid assembly

A hybrid pre-assembly was generated using the genome assembler SPAdes that can combine long PacBio with short Illumina reads. The input consisted of all corrected and filtered PacBio reads together with 60% randomly extracted Illumina reads (321 Million read, 96.3 Gb, to decrease assembly run time and memory requirements). The pre-assembly consisted of 170,143 contigs with a total length of 176 Mb and an N50 of 6,446 bp, of which only 21,690 contigs were larger than 1 kb. CAT filtering was applied to the contigs of the pre-assembly, CAT marked 16,518 contigs consisting of 91.5 Mb (52% of total assembled bases) as contaminant sequences. Next, Illumina reads were aligned to these and Illumina read-pairs of which at least one end aligned were removed from the data set. A final assembly was generated with the CAT-filtered PacBio and remaining 77.6 million Illumina reads, resulting in 8,635 scaffolds with a total length of 32.4 Mb. The assembly size corresponds with the estimate genome size of 36,18 Mb that was determined based on *k*-mer count frequency (Table 1) in the filtered Illumina reads.

**Table 1.** Summary of statistics for the hybrid assembly of the *Pfs1* genome. Data is provided for the final assembly (*Pfs1* final) and size-filtered assembly omitting the scaffold smaller than 1 kb (*Pfs1* filtered). In addition, genome information based on *k*-mer counting of the Illumina reads is providing an estimate for the predicted genome size and repeat content.

|  | *Pfs1* filtered |  | *Pfs1* final |
| --- | --- | --- | --- |
| Assembly size | 30.48 Mb | Assembly size | 32.40 Mb |
| GC content | 47.80% | GC content | 47.75% |
| Longest scaffold | 310.10 kb | Longest scaffold | 310.10 kb |
| Repeat size | 6.38 Mb | Repeat size | 6.93 Mb |
| # Contigs | 3,608 | # Contigs | 8,635 |
| N50 | 36,273 bp | N50 | 32,837 bp |
| # Gene models | 9007 | # Gene models | 13,227 |

|  | **k-mer estimation** |
| --- | --- |
| Assembly size | 36.18 Mb |
| Repeatsize | 8.76 Mb |
| Read Error Rate | 1.04% |
| Model Fit | 94.01% |

### Filtering results

The effect of filtering with CAT on the pre-assembly is well visualized by plotting the GC-content of the contigs, similar as for the PacBio reads. As is visible in Fig 4a many contigs with a GC-percentage deviating from the 40-55% range are present in the pre-assembly, indicating that it contains many contaminating sequences. After filtering, the final assembly (H7) shows one peak of the expected GC-content of 50%, although there is a small shoulder remaining of slightly higher GC-content.

**Fig 4.**
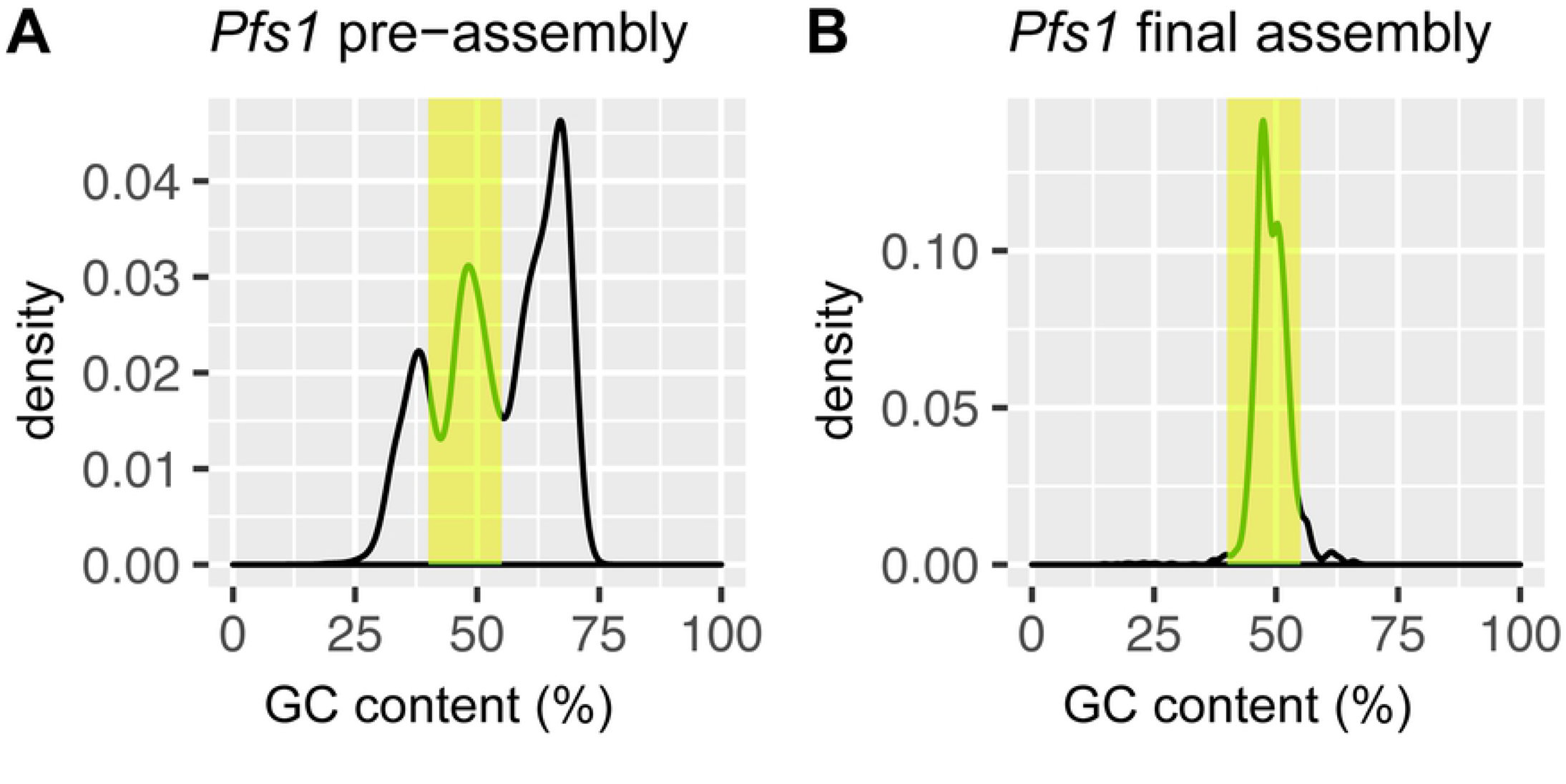
Density plots of the GC values before and after the assembly with filtered reads. Analysis was restricted to contigs > 1 kb. The green bar indicates the region between 45 and 49% GC **A.** GC content of the *Pfs1* contigs from the pre-assembly before filtering shows additional peaks at around 30 and 60 GC%, indicating that there are many contaminant contigs. **B.** GC content of the *Pfs1* contigs after filtering of the reads with the CAT tool shows that the additional peaks are no longer present and have thus been successfully filtered out.

To assess the effectiveness of the taxonomic filtering with a complementary tool we used Kaiju [34]. Kaiju is typically used for the taxonomic classification of sequencing reads in metagenome analysis but here we used it to determine the effect of taxonomic filtering by CAT. For this, genome assemblies of *Pfs1* and other oomycetes were divided into artificial short reads. The taxonomic distributions generated by Kaiju provide a clear picture of the removal of contaminating sequences from the *Pfs1* genome data (Fig 5). Whereas the pre-assembly mostly contained artificial reads with an assigned bacterial taxonomy, this was reduced to 14% in the final assembly. The percentage of >80% of oomycete-assigned reads in the *Pfs1* final assembly is similar to what we observe for the high-quality genome assemblies of *P. infestans* and *P. sojae*, pathogens that can be grown axenically, i.e. free of contaminating other microbes (Fig 5).

**Fig 5.**
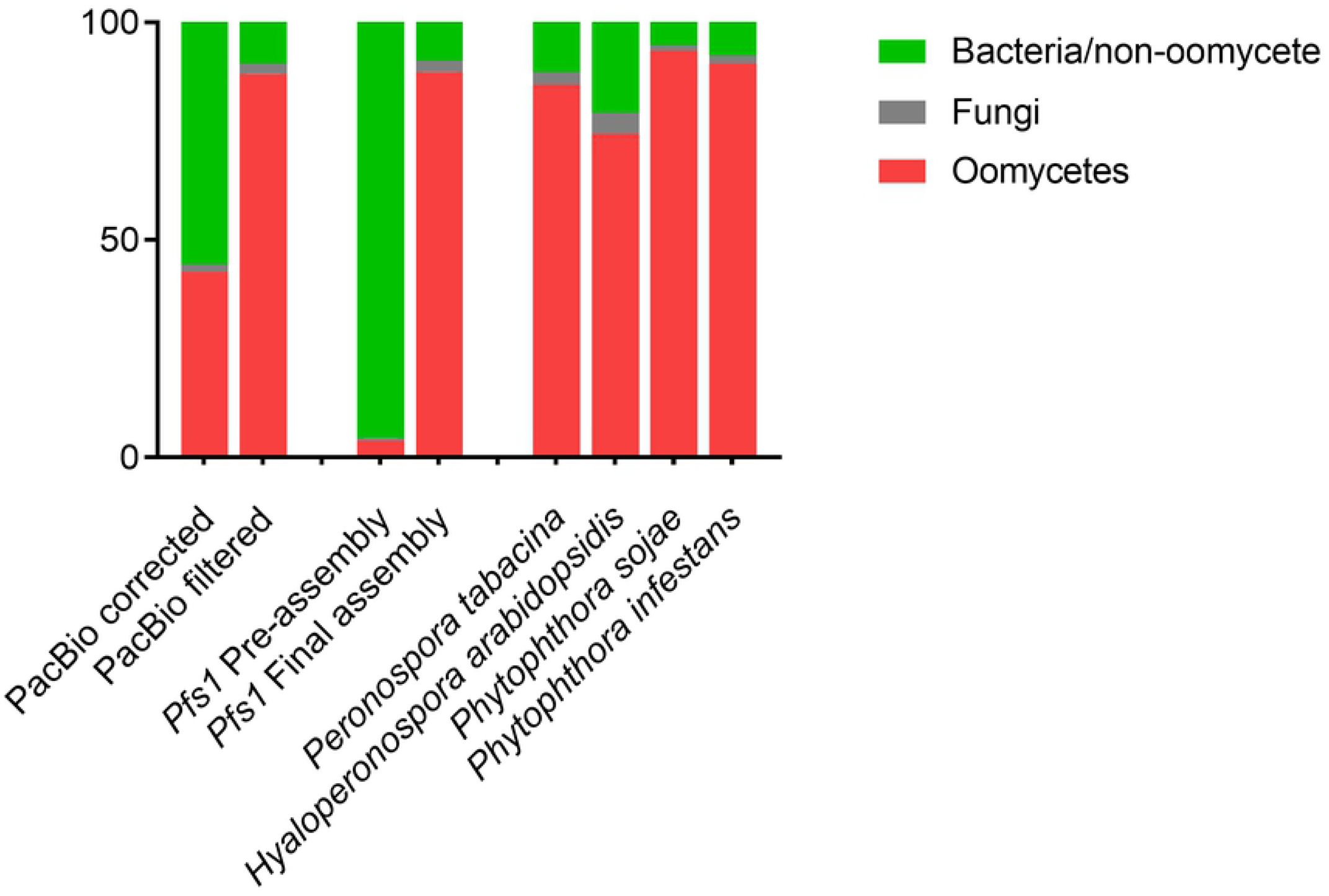
Taxonomic classification of reads in assemblies of different oomycetes. Kaiju bar plot showing the percentage of reads assigned to three taxonomical classes; Oomycetes, Fungi and Bacteria and other non-oomycetes. In error corrected PacBio reads 42.64% are assigned to oomycetes, after filtering with CAT 88.09% of the reads are assigned to oomycetes. For the pre-assembly, only 5% of the reads is assigned to oomycetes. For the *Pfs1* final assembly (H7), 88.6 % of the reads are assigned to oomycetes. This is comparable to other oomycetes that can be sterile grown on plates, indicating that the remaining non-oomycetes are most likely a result of an incorrect classification in the database.

### Genome statistics

To assess the quality of the assembly we re-aligned the Illumina reads to the contigs and found a large variation in coverage between the contigs smaller than 1 kb and the larger contigs, suggesting that these small contigs contain a high number of repeats or assembly errors. In addition, the CAT pipeline depends on classification of individual ORFs on contigs, so it’s accuracy may be expected to improve with contig length. Therefore, several small contigs could possibly be derived from microbes other than *Pfs*. Removing contigs smaller than 1 kb (5027 contigs) resulted in a small reduction of 1.9 Mb in genome length, slightly reducing the assembly size to 30.5 Mb, but resulting in a 58% reduction in the number of contigs. The remaining 3608 contigs, larger than 1 kb, had an N50 of 36,273 bp. The statistics of the filtered assembly are further detailed in Table 1.

To assess the gene space completeness of our assembly in comparison to other oomycete genomes we used BUSCO that identifies single core orthologs that are conserved in a certain lineage. Here, we used the protist Ensembl database as the protist lineage encompasses the oomycetes and other Stramenipila. According to the BUSCO analysis the gene space in our assembly is 88.9% complete with only 0.5% fragmented genes and 0.5% duplicates. This gene space completeness score is similar to that of other downy mildew genomes, but slightly lower than of genomes of *Phytophthora* species (S2 Table). Furthermore, the low number of duplicates suggests that there is a low incidence of erroneous assembly of haplotypes, therefore we concluded that the obtained *Pfs* assembly represents most of the single-copy gene portion of the *Pfs* genome [36].

### Repeat content

In addition to a genome size estimate, the *k*-mer analysis estimated a repeat content of ∼8.8 Mb. This is slightly higher than the observed repeat content in the final assembly of ∼6.9 Mb (∼6.4 Mb in the filtered assembly) (Table 1). The difference between the estimated repeat size and the repeat content in the assembly (1.87 Mb) is most likely caused by long repetitive elements that are hard to assemble. Repeatmasker [20] identified a total of 13,089 repeat elements of which most are part of the Gypsy and Copia superfamily. We also identified 562 LINEs (Long interspersed nuclear elements) and only 16 SINE (short interspersed nuclear elements), these repeat elements belong to the class I transposon (retrotransposons). Other repeat elements consisted of 2297 simple repeats, 298 Low complexity regions, 391 different types of DNA transposons (Table 2), and several (278) other minor repeat types; full details can be found in S3 Table.

**Table 2.** Total number of major repeat types identified in the *Pfs1* genome assembly.

| Repeat type | Count | Percentage | Length (bp) |
| --- | --- | --- | --- |
| LTR | 9247 | 70,65% | 6532069 |
| LINE | 562 | 4,29% | 201127 |
| Simple repeat | 2297 | 17,55% | 97983 |
| DNA repeats/TE | 391 | 2,99% | 46677 |
| Rolling Circle TE | 97 | 0,74% | 26123 |

When we compare the genome assembly size of *Pfs* (30.5 Mb) to other sequenced oomycete genomes such as those of *P. infestans* (240 Mb), *H. arabidopsidis* (100 Mb), Pl. halstedii (75.3 Mb) or the relatively small genome of *P. tabacina* (63.1 Mb), *Pfs* has a strikingly compact genome (S4 Table). The repeat content (21%) is also low compared to that of other oomycetes, e.g. *P. infestans* (74%), *H. arabidopsidis* (43%), *Pl. halstedii* (39% Mbp) and more comparable to *P. tabacina* (24%).

### *Pfs* gene prediction

#### RNA sequencing

Gene prediction is greatly aided by transcript sequence information. We, therefore, isolated and sequenced mRNA from *Pfs* spores and *Pfs*-infected spinach leaves at several time points during the infection. For this, leaves were harvested daily starting from 3 days post inoculation (dpi) until 7 dpi when sporulation was observed. In addition, mRNA was also isolated from sporangiospores and germlings grown from spores that were incubated in water overnight. The 7 different samples would ensure a broad sampling of transcripts that will facilitate gene identification. Illumina transcript sequences (659 million) were aligned to the assembled *Pfs* genome which resulted in 100 million aligned read pairs. Most of the other reads map to the spinach genome but were no further analyzed.

### Predicted proteins

The aligned transcript read pairs served as input for the BRAKER1 [38] pipeline to generate a *Pfs* specific training set for gene model prediction. The gene prediction model was used to predict 13227 gene models on the *Pfs* genome final assembly. The resulting protein models identified in the assembly of *Pfs1* were annotated using ANNIE [40]. ANNIE provided a putative annotation for 7297 *Pfs* proteins (S5 Table). We found that 12630 protein models reside on contigs larger than 1 kb and are thus contained in the filtered assembly. In addition, we found that 2983 gene models had 20% or more overlap with a repeat that was identified by RepeatMasker [20], another 952 protein models were annotated by ANNIE as transposable elements (Fig 6).

**Fig 6.**
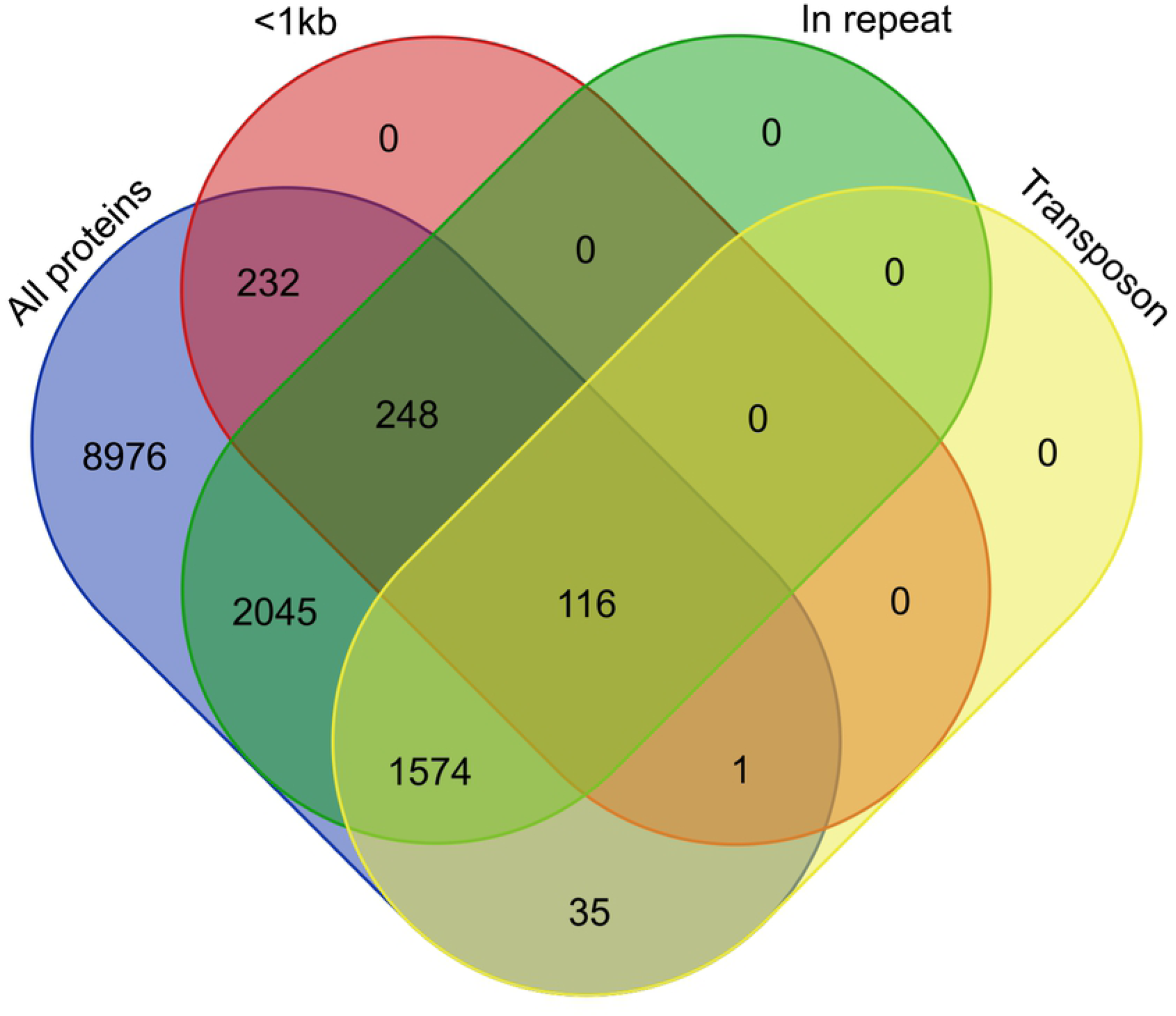
Genomic distribution of protein coding genes. Venn diagram shows the proportion of protein coding genes on contigs smaller than 1 kb (<1k), models that had more than 20% overlap with an identified repeat region in the genome (in repeat) and models that were annotated to be transposons by ANNIE (transposons).

Based on the numbers presented in Fig 7 we conclude that most of the protein models that reside on small contigs (<1 kb) and have a significant overlap with a repeat region are marked by ANNIE as transposons. The number of gene models found in the assembly of *Pfs1* is strikingly low in comparison to that in *P. infestans* (17,792), *H. arabidopsidis* (14,321), *Pl. halstedii* (15,469) and *P. tabacina* (11,310).

**Fig 7.**
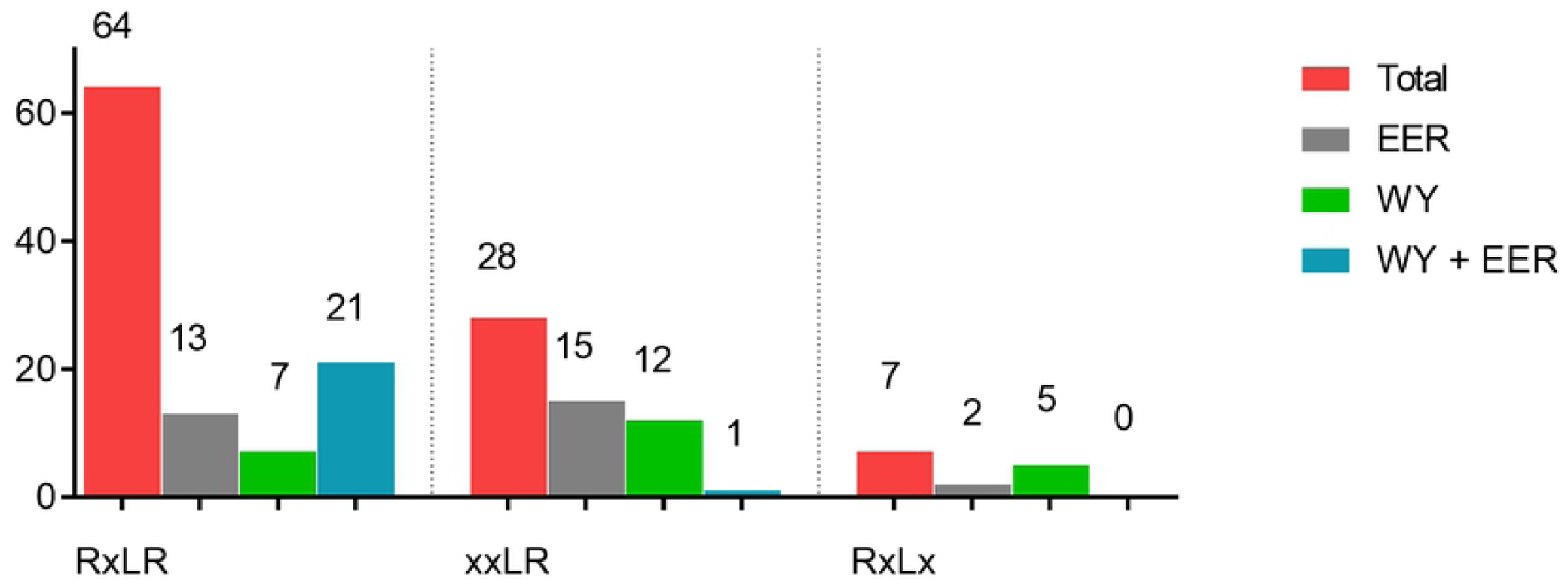
RxLR (-like) motifs observed in the putative RxLR effectors identified in the genome of *Pfs1*. For each (degenerate) RxLR motif the presence of a WY domain (orange), EER-like (green) domain or both (purple) is shown.

### Secretome and host-translocated effectors

For the identification of the *Pfs* secretome as well as of candidate host-translocated RxLR and Crinkler effectors we choose to start with the proteins encoded by the initial 13,227 gene set. This reduces the risk of missing effectors that are encoded on smaller contigs (< 1 kb). SignalP [42] prediction identified 783 proteins with a N-terminal signal peptide. Of these, 231 were found to have an additional transmembrane domain (as determined by TMHMM [44] analysis) leaving 557 proteins. In addition, five of these carried a C-terminal H/KDEL motif that functions as an ER retention signal. The resulting set of 552 secreted proteins was used for secretome comparison, which is ∼ 4% of the *Pfs1* proteome.

Previous research showed that some effectors of the lettuce downy mildew species *Bremia* have a single transmembrane domain in addition to the signal peptide [64]. Therefore, we chose to predict the host-translocated effectors not only from the secretome but also from the set of proteins with a signal peptide and an additional transmembrane domain. A total of 99 putative RxLR or RXLR-like proteins and 14 putative Crinkler effectors were identified. Ten putative RxLR effector proteins were found to have a single transmembrane domain. Also, five putative RxLR effectors are located on contigs smaller than 1 kb (Fig 8, S6 Table). Of the 99 RxLR effectors, 64 had a canonical RxLR domain, while 35 had a degenerative RxLR domain combined with an EER-like and/or WY domain (Fig 7). The number of host-translocated effectors in *Pfs* is significantly smaller compared to that of *Phytophthora* species. Five of the identified putative Crinkler effectors had a canonical LFLAK domain. The others had a degenerative LFLAK combined with an HVL domain or were identified using the Crinkler HMM (Fig 8).

**Fig 8.**
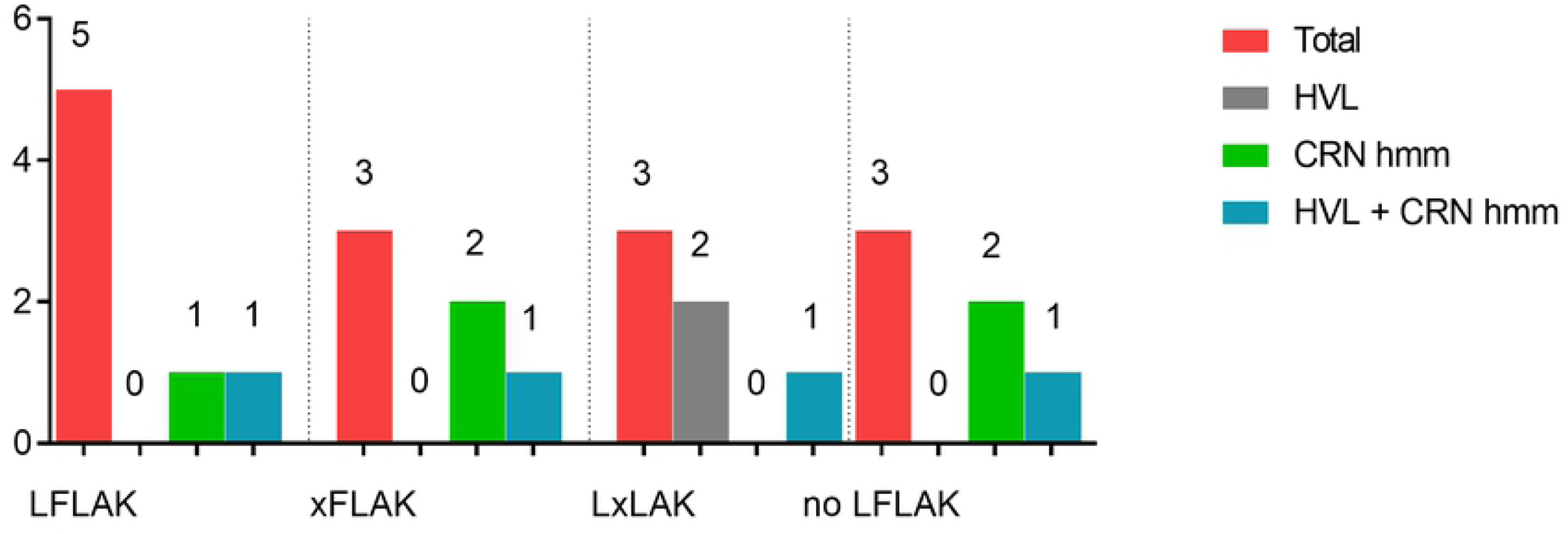
CRN (-like) motifs observed in the putative CRN effectors identified in the genome of *Pfs1*. For each (degenerate) CRN protein the presence of an HVL domain (orange), identified with an CRN HMM model (red) or both (green).

### Genomic distribution of effectors

It has previously been described for the potato late blight pathogen *P. infestans* that effectors often reside on contigs with a relatively large repeat content compared the rest of genome [65]. The distance between neighboring genes was measured to get an idea of the genomic context of the 13277 *Pfs1* genes in general and for 66 selected RxLR effector (canonical RxlR and degenerative RxlR with WY-motifs) genes specifically. To get a good overview of the gene intergenic distances in the genome we plotted the 3’ and 5’ values for all the genes in the *Pfs1* genome on a log10 scaled heat map (Fig 9).

**Fig 9.**
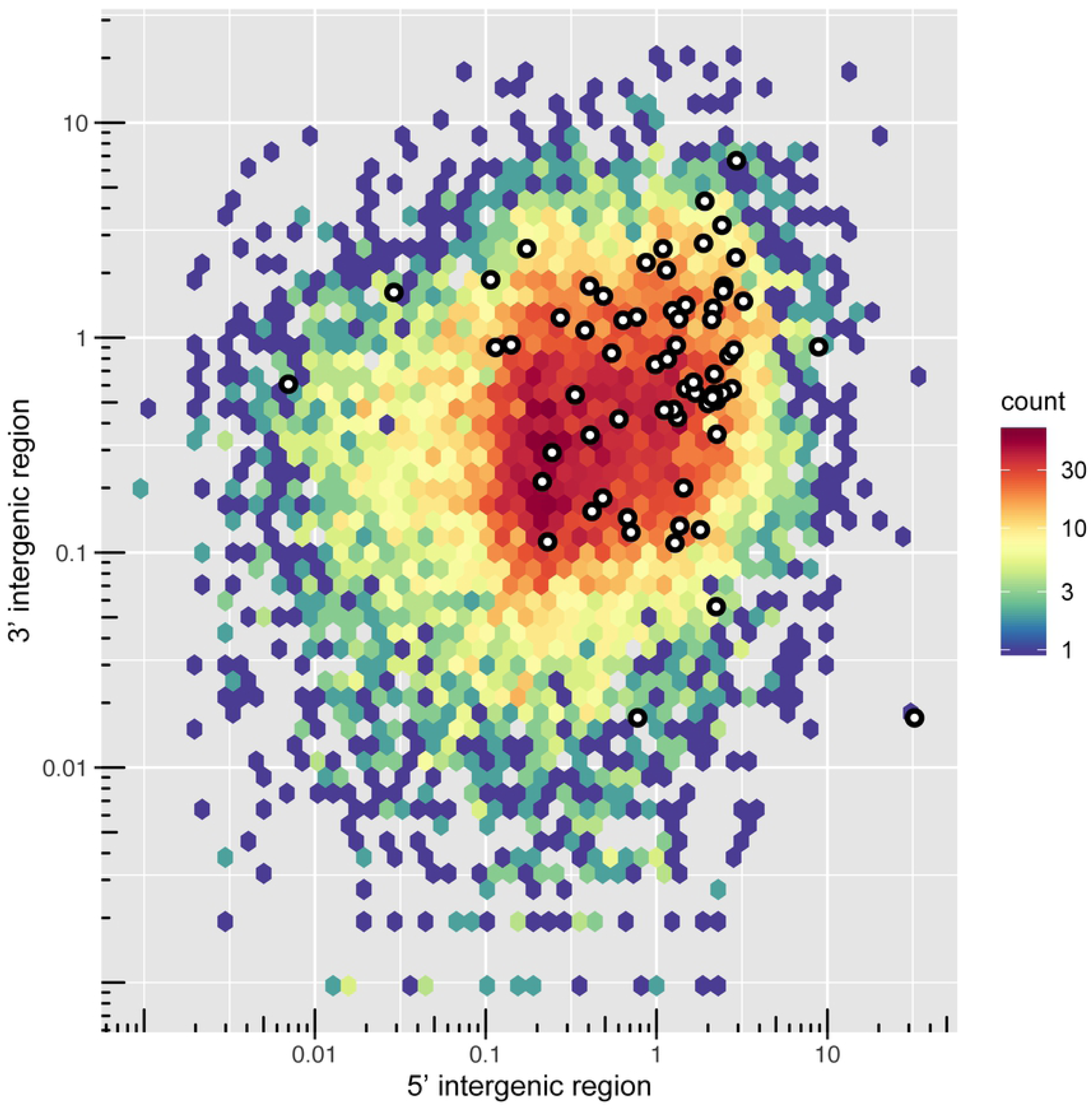
Genome spacing of predicted genes of *Pfs1*. The distance between neighboring genes was depicted by plotting the 5′ and 3′ intergenic distances (on a log10 scale) for each if the 13,227 predicted genes. The scale bar represents the number of genes in each bin, shown as a color-coded hexagonal heat map in which red indicates a gene dense and bleu a gene-poor region. The locations of effectors genes are indicated with white dots. Putative effector genes are plotted in white.

The genome of *Pfs1* is highly gene dense and effectors show a modest but significant (Wilcoxon rank sum test, p = 1.914e^-11^) enrichment in the gene-spare regions of the genome (Fig 9). The median 3’ and 5’ spacing for all genes is 925 bp, while for the selected effector genes it is 2976 bp. However, the difference in gene density between the effectors and core genes is not as strong as in the *P. infestans* two-speed genome [29].

### Comparative analysis of orthologs

Eighteen phytopathogenic oomycete species, that represent a diverse taxonomic range and different lifestyles, were chosen for a comparative analysis with *Pfs* (Table 3). The objective of the comparison is to see whether the biotrophic lifestyle of downy mildew species, like *Pfs*, is reflected in the secretome. For the analysis, the secretome of *Pfs* was compared to that of closely related *Phytophthora* (hemibiotrophic), *Plasmopara* (biotrophic) and more distantly related *Pythium* (necrotrophic) and *Albugo* (biotrophic) species. First, the predicted proteins of each species were used to create a multigene phylogenetic tree to infer their taxonomic relationships using Orthofinder. In total, 86.9% (267,813) of all proteins were assigned to 14,484 orthogroups. Of those, 2383 had proteins from all species in the dataset of which 152 groups contained only one copy of a gene from each species. These single-copy orthologous genes of each species were used to infer a Maximum-likelihood species tree (Fig 10).

**Fig 10.**
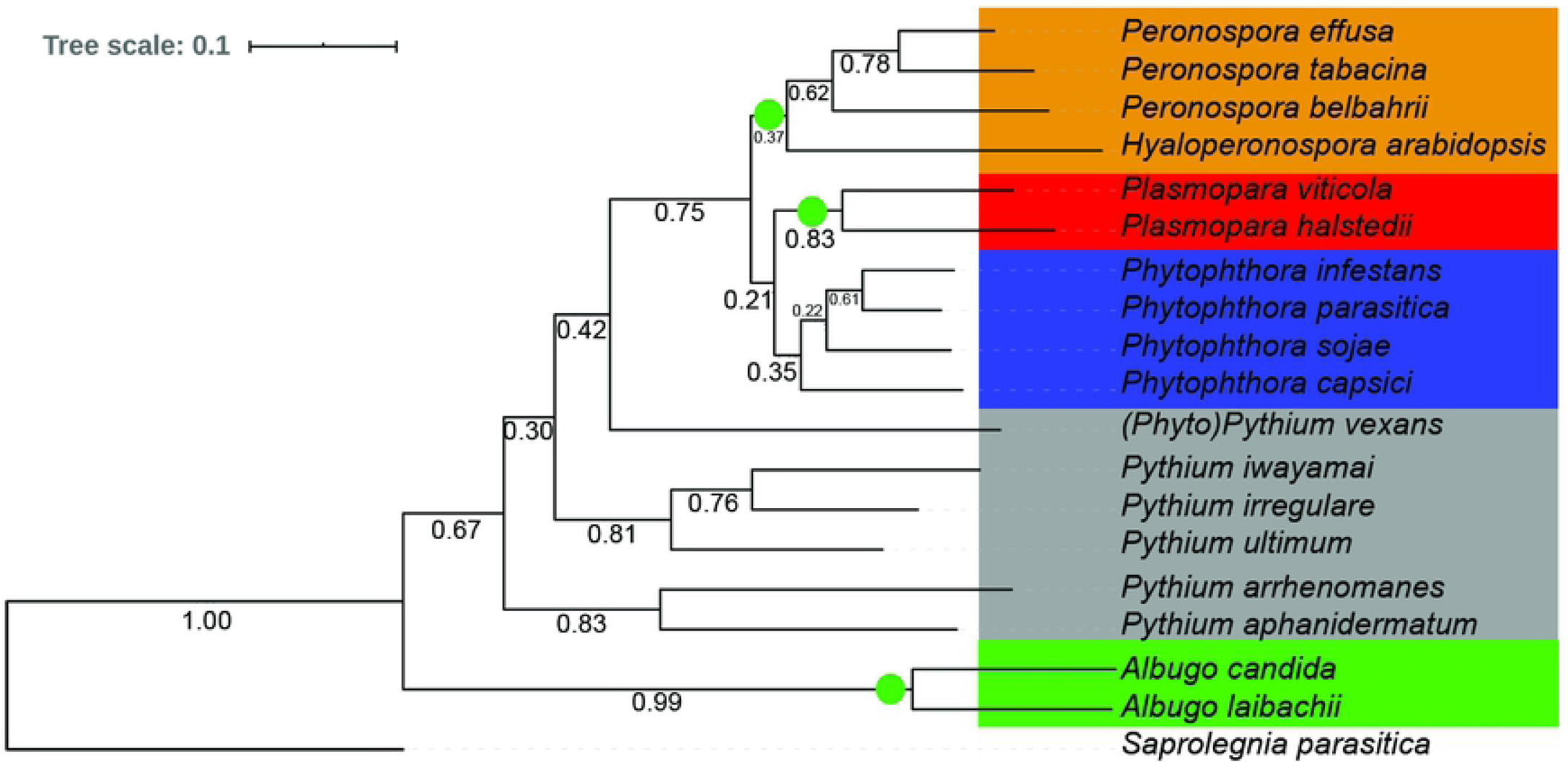
Maximum likelihood tree of the proteome of 18 plant infecting oomycete species. The tree was inferred from 152 single copy ortholog groups in which all species in the comparison where represented. Branch numbers represent bootstrap values of N=12171 trees. Five taxonomic clusters were defined for further analysis;

**Table 3.**
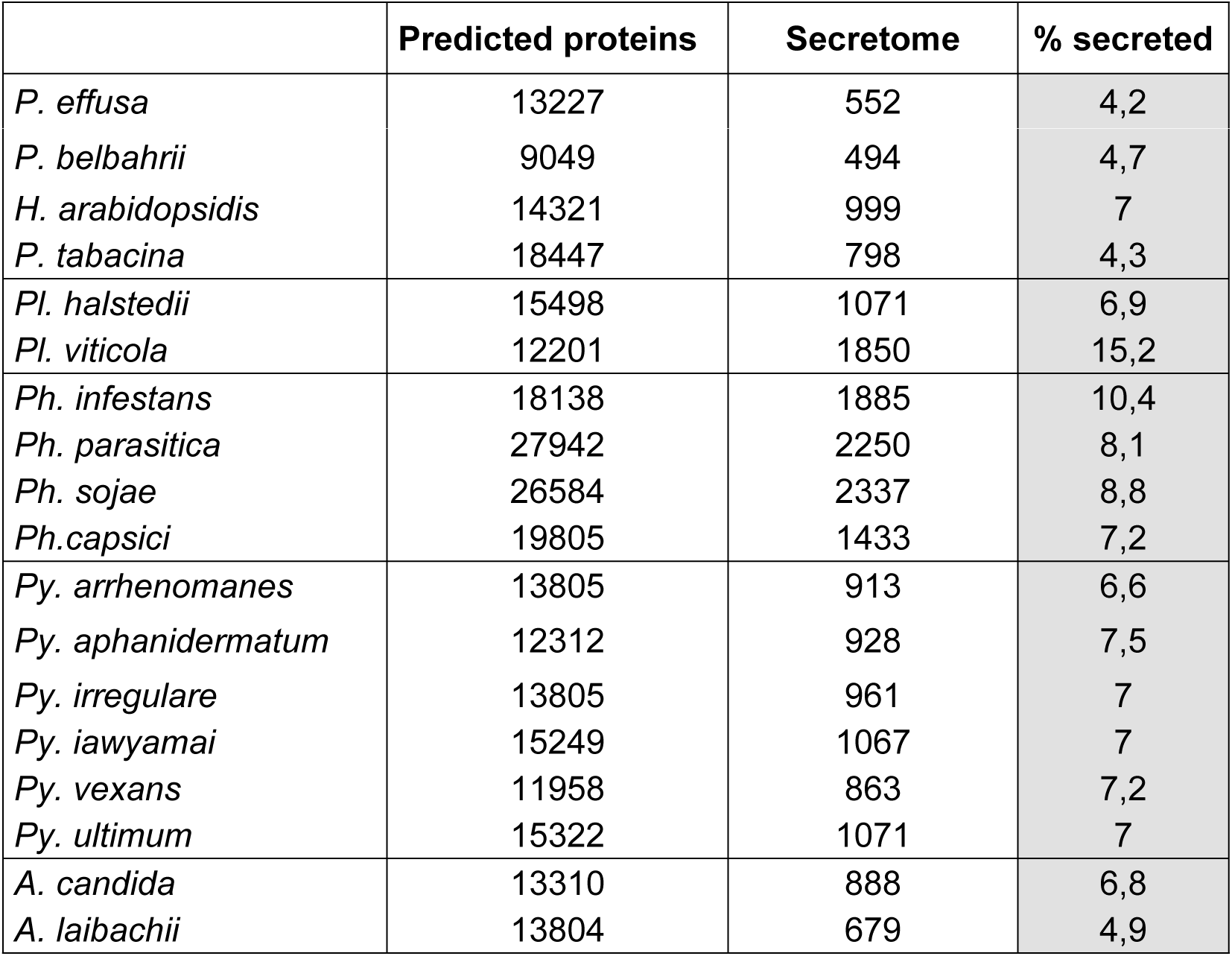
Predicted secretomes of 18 oomycete species used in this study. The total number of predicted proteins, those with a signal peptide (SP), proteins with SP but without additional transmembrane domains (TM), and the number of proteins with SP, no TM, and no C-terminal KDEL sequence are shown. In the final column the percentage of the proteome that is predicted to be secreted is highlighted.

The resulting tree shows that *Pfs* clusters with *H. arabidopsidis* (*Hpa*), *P. tabacina* (*Pta*) and *P. belbahrii* (*Pbe*). The closest relative of *Pfs*, in this study, based on single-copy orthologs is the downy mildew of tobacco *Pta*, followed by the basil-infecting *Pbe*. Based on the tree, *Hpa* is more divergent from the former three downy mildew species within the *Hyaloperonospora/Peronospora* clade. The *Plasmopara* downy mildew species are in a different clade that is more closely related to the *Phytophthora* species used in this study. The separation between the *Peronospora* lineage and the *Phytophthora*/*Plasmopara* lineages is well supported with a bootstrap value of 0.75. This clustering pattern is in line with the recent studies that suggest that the downy mildew species are not monophyletic within the Peronosporales [2, 66]. The *Phytophthora* species, although belonging to three different *Phytophthora* clades, are more closely related to each other than to the other species in this study. *Phytopythium vexans* appears as a sister group to the *Phytophthora/Peronospora* lineage, which is in line with a recently published multi gene phylogeny [67]. The other five species of *Pythium* form two clusters, as previously observed [67]. The two *Albugo* species form a cluster that is separated from the other clades with maximum bootstrap support.

Based on the core ortholog protein tree, we grouped the species into five phylogenetically-related clades; *Hyaloperonospora/Peronospora, Plasmopara, Phytophthora, Pythium* and *Albugo* for further analysis of the secretome. Three of these clades have obligate biotrophic species (*Hyaloperonospora/Peronospora, Plasmopara* and *Albugo*), the *Phytophthora* cluster is hemibiotrophic and the species in the *Pythium* cluster have a necrotrophic lifestyle. (*Phyto)Pythium vexans* was included in the *Pythium* cluster. The fish-infecting *Saprolegnia parasitica* served as an outgroup for the phylogenetic tree and is not used for further comparison.

*Hyaloperonospora/Peronospora* (green), *Plasmopara* (red), *Phytophthora* (blue), *Pythium* (grey) and *Albugo* (green). The obligate biotrophic clades are highlighted using green circle.The fish infecting species *Saprolegnia parasitica,* was used as an outgroup.

### Secretome comparison

For each species, the total number of proteins and the subset that is predicted to be secreted (signal peptide, no additional transmembrane domains, no ER retention signal) is shown in Table 3. *Phytophthora* species generally have a larger proteome than downy mildew species and secrete a larger percentage of the predicted proteins. The *Phytophthora* species in this study are predicted to secrete 1976 proteins on average, whereas the *Plasmopara* and *Peronospora* species secrete an average of 1461 and 703 proteins, respectively.

### Carbohydrate active enzymes

To functionally compare the secretomes between the five defined clades of species we first investigated the carbohydrate-active enzymes (CAZymes). These enzymes, among others, are involved in degrading and modifying plant cell walls, which is an important part of the infection process. Degradation and modification of cell walls is mediated by large numbers of enzymes that have complex interactions. A total of 95 unique CAZyme domains were found, using the CAZymes database (via dbCAN2.0 [55]), in the combined secretomes of the 18 oomycete species. The total number of CAZymes per species ranges from 35 in *A. laibachii* to 336 in *P. sojae*, and was lower in obligate biotrophic species (35 - 193) compared to *Phytophthora* species (197 - 336) (S7 Table).

The presence and numbers of CAZyme domains were compared between species using a Principal Component Analysis (PCA), a statistical reduction technique that determines what variables contribute most to the variation observed in a data set. We report the relative abundance of each CAZyme domain to the total number of secreted CAZyme domains per species, to account for the large variation in absolute numbers of proteins between the species (Fig 11). A PCA based on the absolute numbers can be found in S2 Fig, which shows a similar pattern. The species clusters as depicted in Fig 11 are supported by PERMANOVA (p < 0.001).

**Fig 11.**
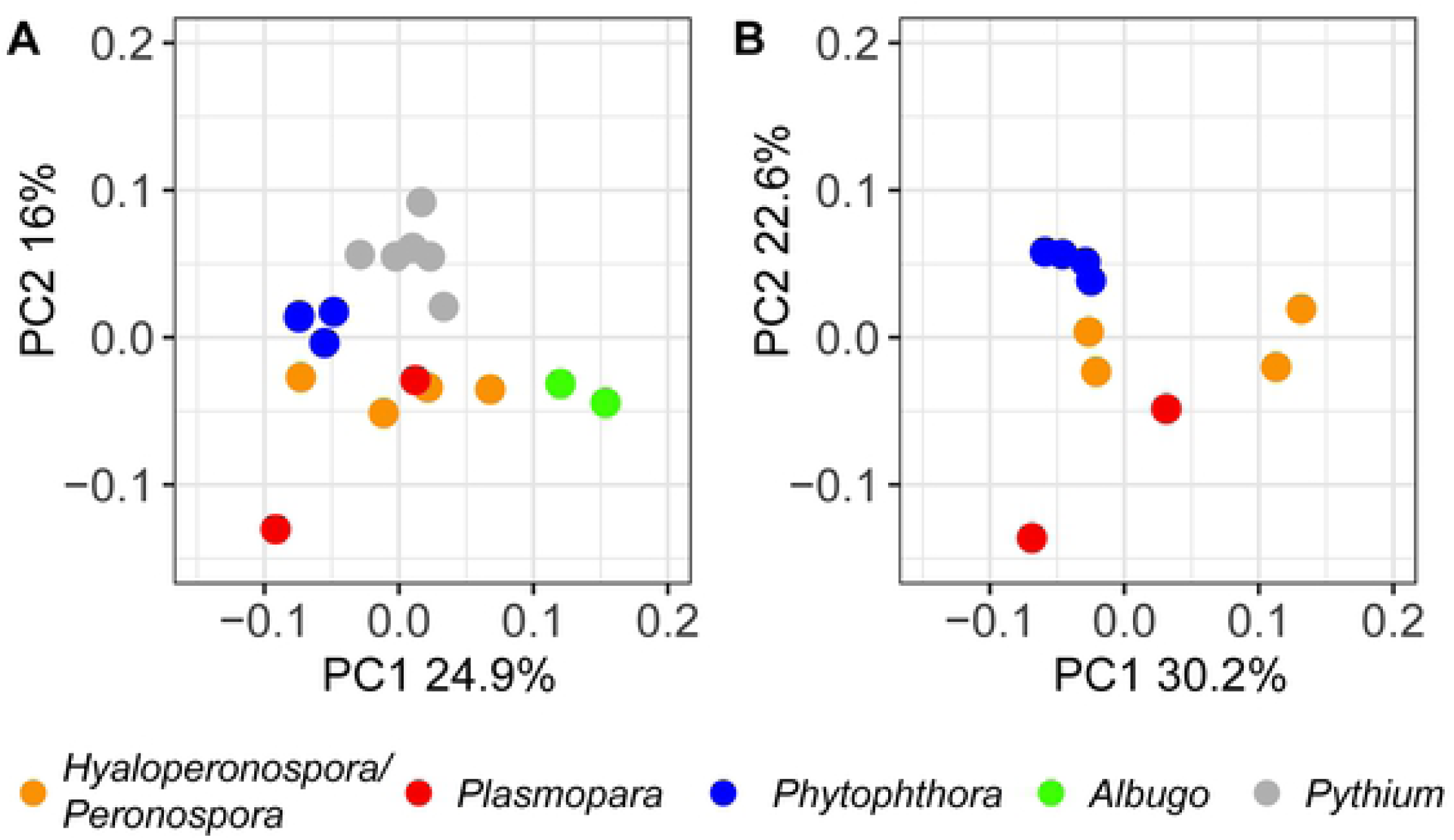
Principal component analysis (PCA) of variation in the relative abundance of secreted CAZymes. PCA was done for all of the 18 species (**A**) or for the *Peronospora*, *Plasmopara* and *Phytophthora* species only (**B**). The PERMANOVA test shows that the grouping based on the CAZyme domains is significant (Pr>F = 0.001). Species are grouped by color based on the classes that where defined in the phylogenetic tree (Fig 10). *Phytophthora* (blue), *Peronospora* (yellow), *Plasmopara* (red), *Albugo* (green) and *Pythium* (grey). Abbr. *PFS; Peronospora (P.) effusa, PBE; P. belbahrii, PTA; P. tabacina, HPA, Hyaloperonospora arabidopsidis, PHA; Plasmopara (Pl.) halstedii, PVI; Pl. vitiocola, PIN; Phytophthora (Ph.) infestans, PSO; Ph. sojae, PCA; Ph. capsici, PPA; Ph. parasitica, ACA; Albugo (A) candida, ALA; A. laibachii, PUL; Pythium (Py.) ultimum, PAR; Py. arrhenomanes, PAP; Py. Aphanidermatum, PIR; Py. irregulare, PIW; Py. Iawyamai, PVE; Phytopythium vexans*

In accordance with the core ortholog protein tree (Fig 10), the *Albugo*, *Phytophthora* and *Pythium* species form separate clusters in the PCA. Remarkably, the *Hyaloperonospora/Peronospora* species do not cluster together, indicating that the secreted CAZyme domains vary largely between these species, despite their close phylogenetic relationship and same lifestyle. Similarly, the two *Plasmopara* species do not cluster together and are not closer to *Phytophthora* than *Peronospora* as could be expected based on the core ortholog protein tree (Fig 10). Instead, the *Plasmopara* and *Hyaloperonospora/Peronospora* species are not seperated in the PCA based on the secreted CAZyme domains. The repertoire of CAZymes in all these downy mildew species is more similar than would be expected based on their taxonomic relationship. This could be the result of convergent evolution towards the obligate biotrophic lifestyle. However, the *Plasmopara* and *Hyaloperonospora/Peronospora* species are in no way clustered as the position of the individual species is highly variable among component one. The *Albugo* species do not cluster with the other biotrophic species.

To exclude the effect of the more distantly-related species on the separation between the downy mildew and *Phytophthora* species, the PCA was also performed on the set without the *Pythium* and *Albugo* species (Fig 11b). The pattern, as observed in the total set, is maintained when the more distantly related species are excluded from the analysis.

We conclude that a different composition and abundance in secreted CAZyme domains may contribute to obligate biotrophy as these species are clearly separated from the necrotrophic and hemibiotrophic ones. However, the variation within obligate biotrophic species is large making it impossible to find a clear carbohydrate-active enzyme signature that defines the obligate biotrophs.

To look further into the properties of the secreted CAZymes we highlighted literature-curated domains of phytopathogenic oomycetes that are known to modify the main plant cell wall components; lignin, cellulose and hemicellulose [68] (S3 Fig). We found that the secretomes differ more in terms of the absolute number of plant cell wall-degrading enzymes than in the relative occurrence of the different corresponding CAZyme catalytic activities per species. Secretomes of obligate biotrophic and hemibiotrophic/necrotrophic oomycetes have secreted proteins with similar functions (like breakdown of cellulose, pectin, hemicellulose etc.) but the numbers and diversity of those proteins in obligate biotrophic species are reduced.

### Pfam domains

In a different approach the secretomes were compared based on their Pfam annotation. InterPro was used to identify known protein domains and the numbers and presence of domains was compared between the species. A total of 1354 unique domains were found in the combined secretomes of the oomycetes analyzed. The number of domains identified ranged from 304 in *Al. candida* to 1710 in *Ph. parasitica.* The total number as well as the relative number of Pfam domains in secretomes of obligate biotrophic species was lower in obligate biotrophic species compared to *Phytophthora* and *Pythium* (S8 Table).

PCA analysis was used to compare the Pfam annotations of the secretomes based on relative number of secreted Pfam domains per species (Fig 12a, absolute numbers S4 Fig). The PCA shows a clear separation between lifestyles. The *Phytophthora* species cluster together and separate from all other species along PC1 (25,3%). The *Pythium* species form a cluster that separates clearly from the other species along PC2 (20,2%). Remarkably, all biotrophic species, including both groups of downy mildews and the *Albugo* species cluster together. Within the obligate biotrophic cluster the phylogenetic groups (*Hyaloperonospora/Peronospora, Plasmopara, Albugo)* as found in the core ortholog tree are still present but the differences are minor. The species clusters as depicted in Fig 12 are supported by PERMANOVA (p = 1e^-3^). To exclude the effect of the more distantly related species on the separation between the obligate biotrophs, the PCA was also performed without *Pythium* and *Albugo* species (Fig 12.b). The pattern observed in Fig 12a is maintained when the more distantly related species are excluded from the analysis (Fig 12.b).

**Fig 12.**
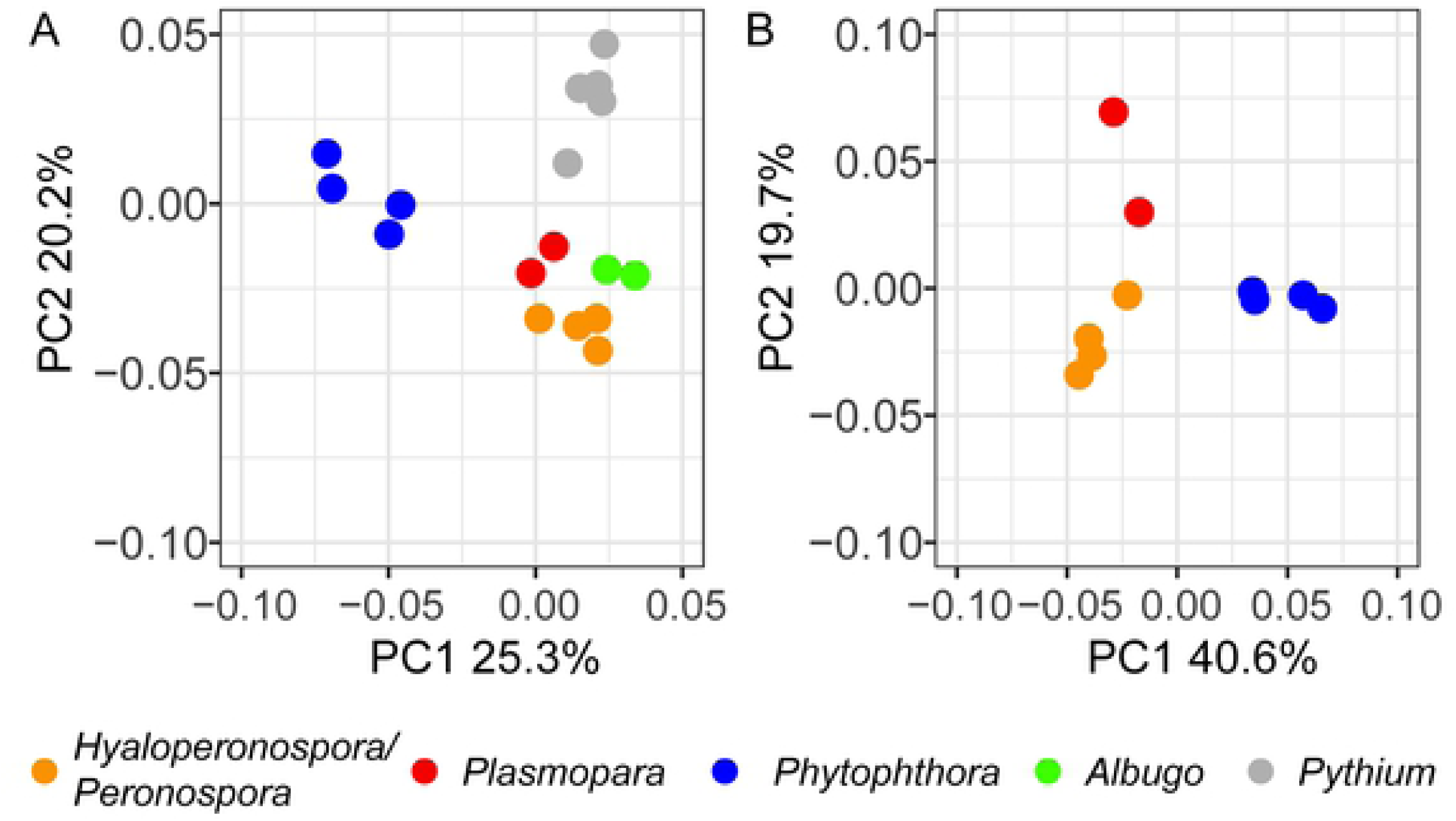
Principal component analysis (PCA) of variation of the relative abundance of predicted secreted Pfam domains. In (**A**) all species are included, while (**B**) shows the variation between downy mildew, *Plasmopara* and *Phytophthora* species only. Species are grouped by color based on the classes that where defined in the phylogenetic tree (Fig 10). The PERMANOVA test shows that the grouping based on the Pfam domains is significant (P < 0.001). *Phytophthora* (blue), downy mildew (yellow), *Plasmopara* (red), *Albugo* (green) and *Pythium* (grey).

The Pfam domains that contribute to the variance in PC1 and PC2 were identified using a biplot. In a biplot, the variables are presented as vectors, with their length reflecting their contribution. Many of the domains contribute to the differences between the biological groups, but five of them stand out (Fig 13. and Table 4).

**Fig 13.**
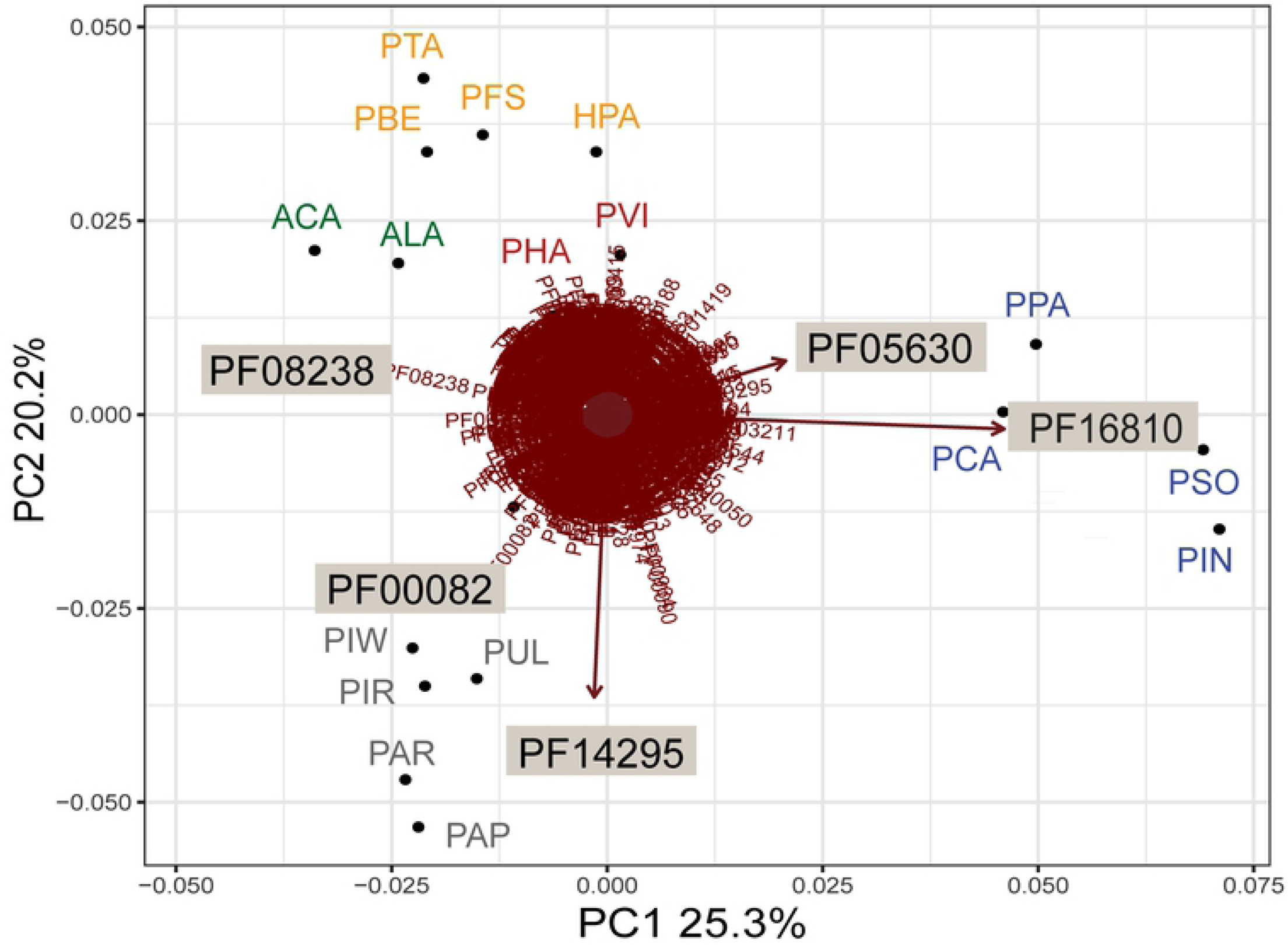
Pfam domains that provide to the variation in the relative abundance between species. Although many domains contribute to the variation, PF16810, PF05630, PF08238, PF14295 and PF00082 are the domains that contribute most, as evidenced by the length of their vectors in the biplot.

**Table 4.**
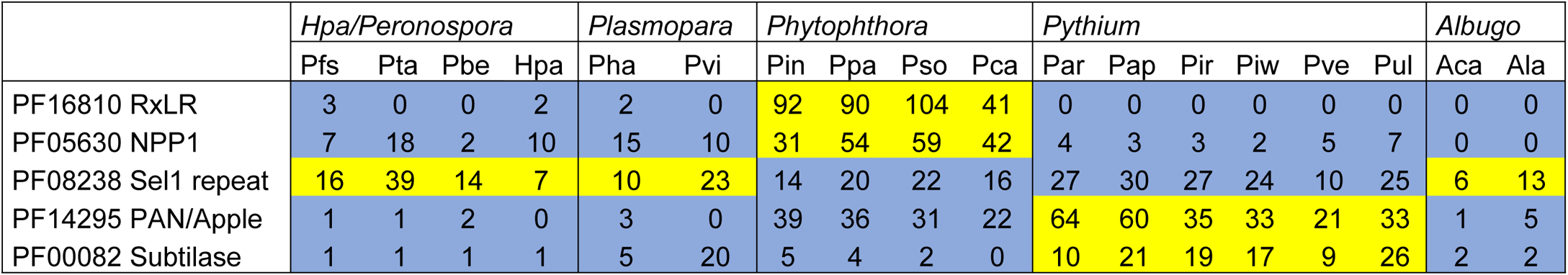
Pfam domains that contribute most to the variation between species in the PCA. Numbers represent the number of domains per secretome per species. Domains that are relatively less abundant are blue, domains that occur in relatively high numbers are yellow.

Two Pfam domains that have a higher relative abundance in *Phytophthora* contribute strongly to the separation between *Phytophthora* and the other species. The first, PF16810, represents a RxLR protein family with a conserved core α-helical fold (WY-fold). Some of the proteins that this domain was based on have a known avirulence activity [50]. On average, 82 PF16810 domains were identified in *Phytophthora* compared to 1.3 in *Peronospora*, 1.0 in *Plasmopara* and none in the *Albugo* clade. Using HMMer searches, many more WY domain candidates can be identified in these species, however they apparently do not fit the Pfam domain that is based on a larger part of the proteins as the HMM.

The second, PF05630, is a necrosis-inducing protein domain (NPP1) that is based on a protein of *Ph. parasitica* [69]. This domain is conserved in proteins belonging to the family of Nep1-like proteins (NLPs) that occur in bacteria, fungi and oomycetes [70]. Infiltration of cytotoxic NLPs in eudicot plant species results in cytolysis and cell death, visible as necrosis [71]. *Phytophthora* species are known to have high numbers of recently expanded NLP genes in their genomes, encoding both cytotoxic and non-cytotoxic NLPs [70]. *H. arabidopsidis* and other obligate biotrophs tend to have lower numbers and only encode non-cytotoxic NLPs [70, 72].

Domain PF08238 contributes to the distance between the *Phytophthora* and obligate biotrophic species and is relatively more abundant in the biotrophs (PC1). PF08238 is a Sel1 repeat domain that is found in bacterial as well as eukaryotic species. Proteins with Sel1 repeats are suggested to be involved in protein or carbohydrate recognition and ER-associated protein degradation in eukaryotes [73]. No function of proteins with a PF08238 domain is known for oomycete or fungal pathogens.

The distance between *Pythium* and the obligate biotrophic species along PC2 is largely caused by differences in two domains that are commonly reported in oomycete secretomes [66]. The first, PF14295, a PAN/Apple domain, is known to be associated with carbohydrate-binding module (CBM)-containing proteins that recognize and bind saccharide ligands in *Ph. parasitica*. Loss of these genes, as in the biotrophs, may facilitate the evasion of host recognition as they can induce plant defenses [74]. Second, PF00082, is a subtilase domain, which is found in a family of serine proteases. Secreted serine proteases are ubiquitous in secretomes of plant pathogens and play a role in nutrient acquisition through the breakdown of host proteins [75]. Secreted proteases from fungal species have been shown to enhance infection success by degrading plant derived antimicrobial compounds [76].

### Over and under-representation of Pfam domains in obligate biotrophic species

The enrichment analysis confirmed the pattern that was shown in the biplot. In total, 60 Pfam domains were found to be differentially abundant in obligate biotrophic species clusters compared to *Phytophthora* (Table 5). Three out of the five Pfam domains that contributed most to the separation between phylogenetic groups in the PCA (Fig 12. and Table 4) were also found to be differentially abundant in at least one obligate biotrophic cluster compared to *Phytophthora* in the enrichment analysis. The RXLR domain (PF16810) was under-represented in all three obligate biotrophic clusters. The PAN domain (PF14295) was under-represented in *Plasmopara* and *Peronospora* and the NPP1 domain (PF05630) in *Albugo*. Upon inspection a trend towards under-representation of NPP1 (Bonferroni corrected p = 0.159) in *Hyaloperonospora/Peronospora* is evident as well, although not significantly. The Sel1 domain (PF08238) significantly over represented between *Phytophthora* and *Hyaloperonospora/Peronospora*. The fifth domain that was found to contribute to the variation in the PCA was not found in the enrichment analysis, as it accounts for separation between *Pythium* and the other clusters. No additional domains were found to be differentially abundant in more than one obligate biotrophic species cluster compared to *Phytophthora*.

**Table 5.**
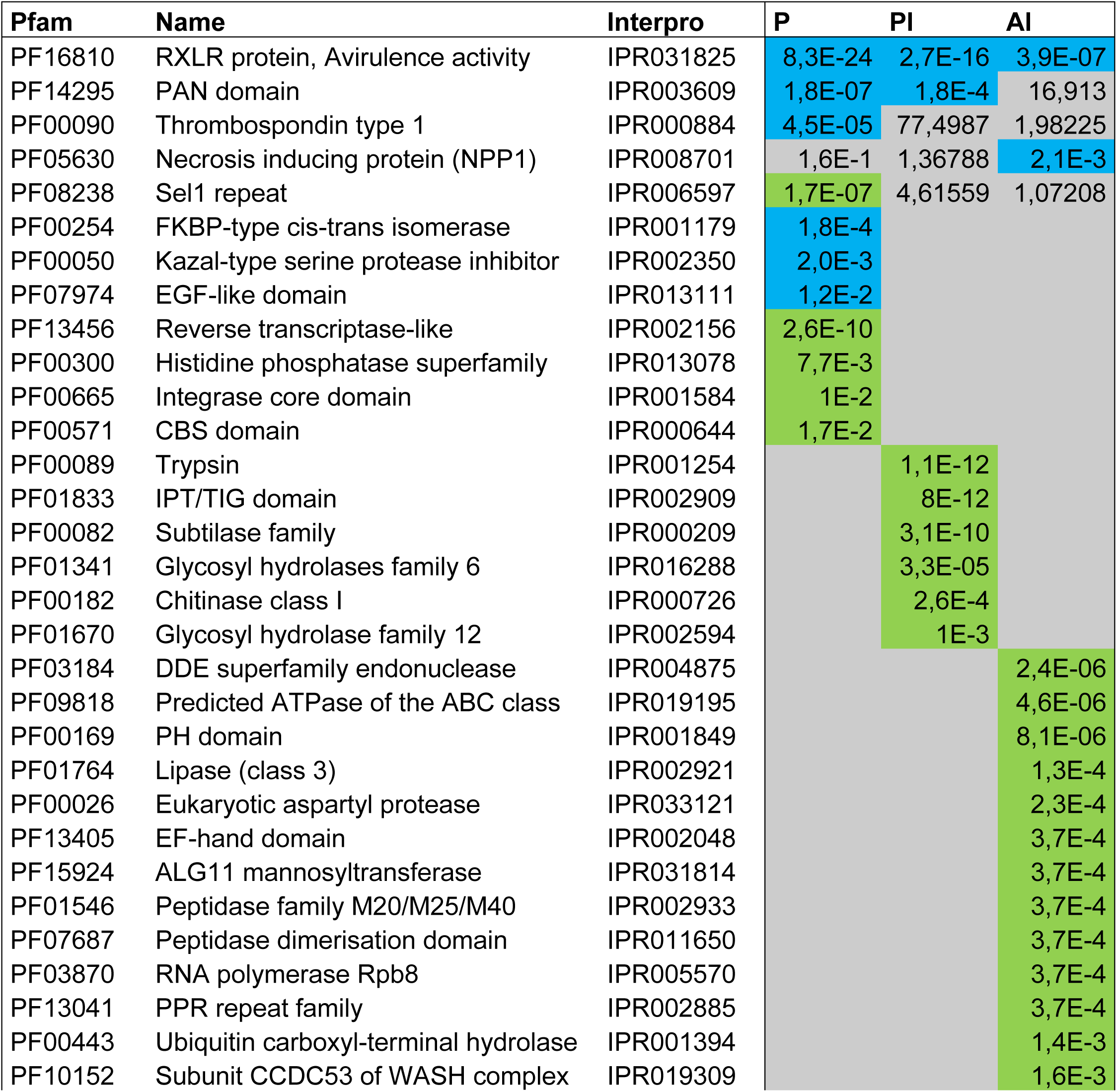
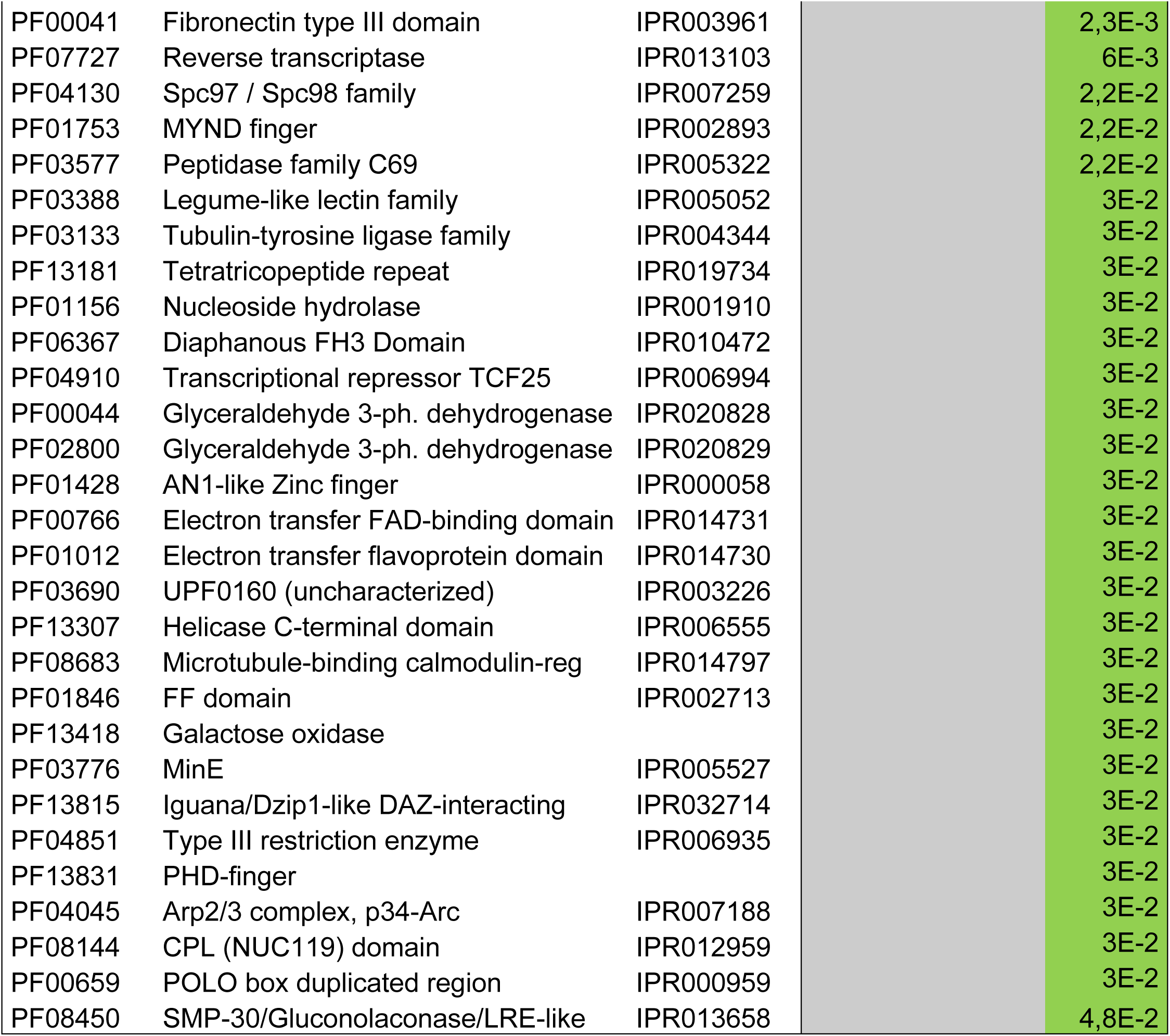
Over and under-representation of Pfam domains in the secretomes of *Hyaloperonospora/Peronospora*, *Plasmopara* and *Albugo* compared to *Phytophthora* species. Over (green) and under (blue)-representation was tested relative to the expected distribution of each Pfam domain. The abundance of each domain was compared between the species clusters using a Chi-square test with Bonferroni correction. Bonferroni corrected p-values are shown in the table.

Previous studies identified Pfam domains that are associated with virulence in other phytopathogenic oomycete species like *Pythium, Plasmopara, Peronospora* and *Phytophthora* [77]. The occurrence of known virulence-associated domains of *Pfs* is summarized in S5 Fig. We found that obligate biotrophic species have a lower total number of secreted virulence-associated domains and that these domains make up a smaller portion of the total number of secreted proteins compared to the other species.

### Host-translocated effectors

The RxLR effector models in the Pfam database (PF16810 and PF16829) mentioned above cover only a small fraction of the predicted RxLR effectors in secretomes of phytopathogenic oomycetes. We predicted the total number of host translocated effectors for each secretome using a Perl regex script and HMM searches (see methods) to get a more comprehensive view of the total numbers of host translocated effectors. This includes RxLR effectors without WY domains and CRN effectors.

The predicted host translocated effectors are classified into groups based on the identified domains and shown in Fig 14. Please note that the number of *Pfs* effectors is slightly different from the numbers reported before. The numbers in Fig 14 are derived from predictions based on published HMM models instead of models trained for *Pfs* (Fig 7), for the purpose of comparison between species. The same criteria for effector prediction were used for all species in this comparison.

**Fig 14.**
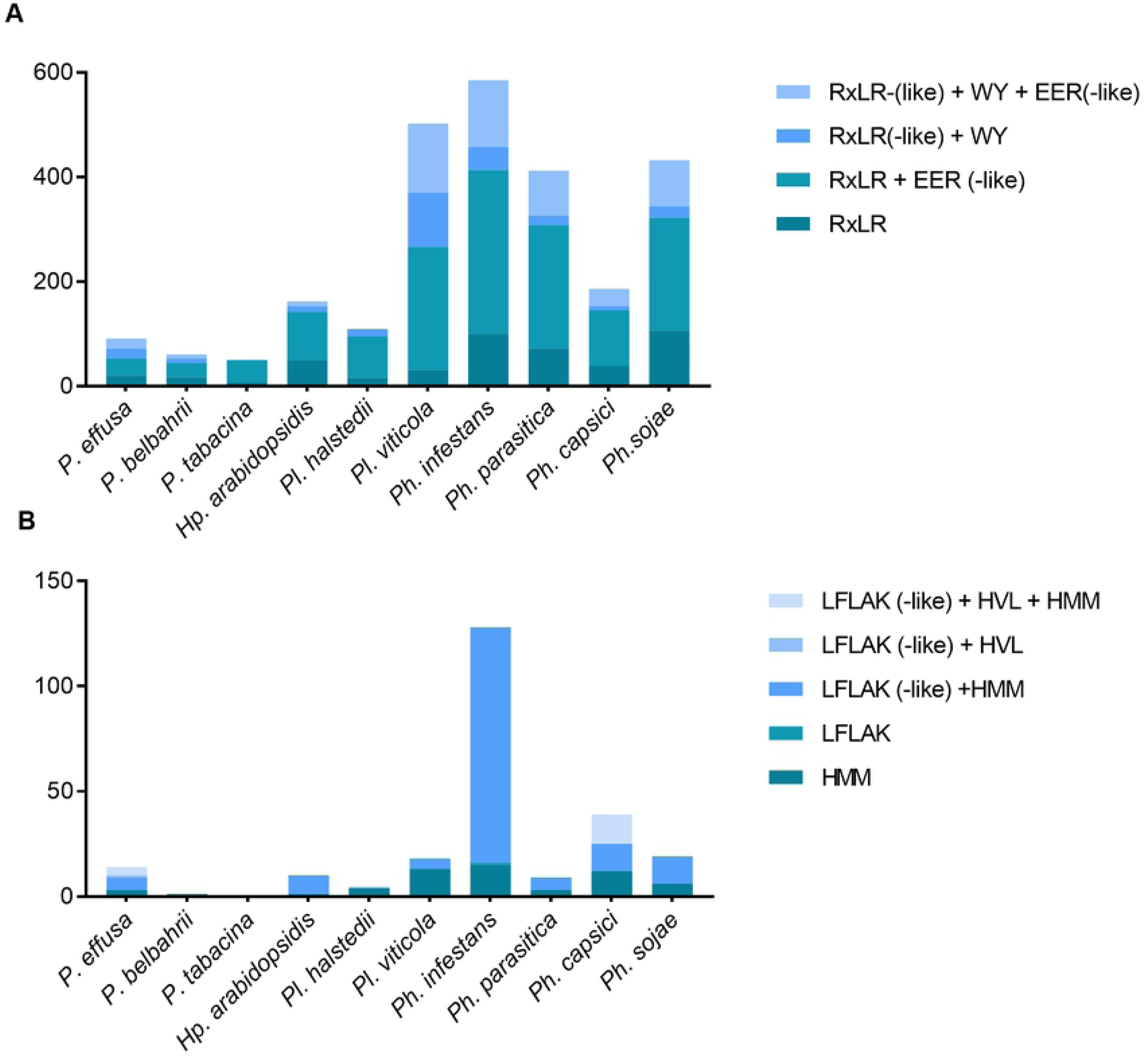
Predicted (a) RxLR and (b) CRN effectors in the secretome of downy mildew, *Plasmopara* and *Phytophthora* species. The predicted effectors are classified into four (RxLR) or five (CRN) categories, based on the domains they possess.

RxLR effector proteins are more abundant in *Phytophthora* compared to the obligate biotrophic species in this study. On average 399 RxLR effector proteins were found in *Phytophthora* whereas *Plasmopara* and *Hyaloperonospora/Peronospora* had 79 and 90 on average. The same pattern is evident for CRN effectors. The average number of CRN proteins in *Hyaloperonospora/Peronospora* is 11, while *Plasmopara* has 12 and *Phytopthora* 56 on average. We conclude that downy mildew species (*Hyaloperonospora/Peronospora* and *Plasmopara*) have fewer host translocated effectors compared to *Phytophthora* species.

## Discussion

### Taxonomic filtering

The ability to sequence full genomes at high pace and relatively low cost has aided research in phytopathology dramatically. Over the past few years the genomes of many phytopathogenic oomycetes have been sequenced and their genomes revealed an arsenal of protein coding genes that were found to be important in virulence. However, technical difficulties restricted the sequencing and assembly of genomes of obligate biotrophic oomycetes that cannot be cultured axenically. Obligate biotrophic species can only grow on living host tissue so when collecting spores for DNA isolation bacterial and plant DNA will inevitably contaminate the sample, which complicates the genome assembly. In this paper we describe the assembly of a relatively clean genome sequence of the obligate biotrophic downy mildew of spinach, *Peronospora effusa*.

To get a clean assembly, sequence that are derived from different species were filtered out and removed. Several methods were considered to identify and filter contigs or reads that were likely contaminants in our data. Initially we considered to filter contigs or reads based on their GC content, since this differs between oomycete genomes [78] and most other microbes [79]. However, not all bacteria have a distinctive nucleotide usage, for example *E. coli* has a GC content of 51.7% [79], which is close to that of most oomycetes. In addition, the GC bases are not evenly distributed over the genome, so filtering based on GC content could therefore potentially remove valuable parts of the genome.

Alternatively, reads of non-oomycete origin could be identified by mapping them to databases with sequences of known taxonomy. For example, a database containing only oomycete or bacterial genomes. This is not ideal as the databases are incomplete and are likely to contain annotation errors. In addition, it could lead to the removal of novel parts of the downy mildew genome that are not present in other oomycetes, eventually this will hamper the study on the valuable species-specific parts of your genome.

The filtering we applied with the CAT tool does not classify a contig based on a single hit. Instead it determines the taxonomic origin of each ORF on an assembled contig or corrected PacBio read, providing a robust classification [12]. After filtering, 50% of the error-corrected PacBio reads remained, and were used in the assembly and 56% of the Illumina sequencing reads aligned with the assembled genome. This indicates that roughly half of our sequenced data originated from another microbial source besides *Pfs*. Notably, while the classifications in the original CAT paper were only benchmarked on prokaryotic sequences [12], our study shows that the tool also performs well for classifying eukaryotic contigs. Thus, the tool may also be promising for classification of eukaryotes including oomycetes in metagenomic datasets, provided that long contigs, or corrected PacBio or Nanopore sequencing reads are available.

It should be noted that sequences of unknown taxonomy were maintained for the assembly, making it possible that these are still contaminants. When we compare the taxonomic distributions generated by Kaiju of the pre-assembly and final assembly, we see a dramatic reduction of sequences of bacterial origin (Fig 5). The oomycete content according to Kaiju and the overall GC content of the final assembly is similar to that of genome assemblies of axenically grown oomycetes. We can therefore conclude that with the CAT filtering method, we successfully filtered out most of the genomic reads that did not belong to the oomycete genus.

### Hybrid assembly

Most oomycete genomes sequenced to date were found to contain long repeat regions [80] that cannot be resolved using only a short read technology such as Illumina. Long reads can potentially sequences over these long stretches of repeat sequences, and contribute to the contiguity of the genome assembly [81]. Therefore, the data was complemented with long read PacBio sequences in an attempt to close the gaps between the contigs. The resulting assembled *Pfs1* genome consist of a large number of contigs. This suggest that even with PacBio reads we were unable to span many repeat regions. Besides biological reason for the large number of contigs, there could also be a technical reason. Prior to PacBio sequencing whole genome amplification with random primers was performed as the initial sequencing attempt with non-amplified DNA barely yield sequencing reads. This creates a bias, where some parts of the oomycete the genome may be differentially represented in the PacBio data set by chance. Nevertheless, CAT effectively removed the bacterial contamination providing a clean input for the assembly.

### The genome of *Pfs1*

The assembled *Pfs1* genome size is 32.40 Mb divided over of 8,635 contigs. The genome is highly gene dense and contains in total 13,227 genes. Overall, the BUSCO analysis showed that this assembly contains most of the gene-space. Many of the 8,635 contigs were smaller than 1 kb. However, the CAT filtering method performs best on relatively large contigs containing multiple ORFs. Therefore, small contigs could still contain sequences derived from other microbes. The removal of these small contigs results in a small genome size reduction (1.9 Mb), but significantly reduces the number of contigs (by 5027). When we account for genes that have a significant overlap (>20%) with repeats, or that were annotated as transposable elements we come to 9,007 high-confidence gene models.

The genomes of *Pfs* race *13* and *14* have recently been published [82, 83], with a similar genome size (32.1 Mb, and 30.8 Mb respectively) and gene content (∼ 8000 gene models) compared to the assembly of *Pfs1*. Contrary to our assembly method, the input data for those genome assemblies were filtered by alignment to an oomycete and bacterial database to discard reads that do not belong to the oomycete genus. This filtering method could potentially lead to the incorporation of bacterial sequences that are not in the public databases. Besides, the positive filtering for oomycete scaffolds against NCBI nt database could have resulted in the loss of *Pfs* specific genome sequences. In addition, the assembly of *Pfs13* and *14* were based on the merging of several assemblies of different *k*-mer characteristics. Although this improves the contiguity of the assembly, this method might yield collapsed and chimeric contigs.

### *Peronospora* species have reduced genomes

Recent sequencing of *Peronospora* species shows that they have remarkably small and compact genomes (32.3 – 63.1 Mb) compared to *Phytophthora* (82-240 Mb) species [29, 33, 82, 84]. The *k-*mer analysis predicts the *Pfs1* genome to be 36.18 Mb containing 8.78 Mb of repeats (24%). The predicted genome size of *Pfs R13* and *R14* based on *k-*mer prediction is 44.1 - 41.2 mb (repeats; 24 – 22%) [82]. The increased genome size of *Phytophthora* is attributed to an ancestral whole genome duplication in the lineage leading to *Phytophthora* and to an increase in the proportion of repetitive non-coding DNA [29, 85]. The duplication event has been proposed to have taken place after the speciation of *Hp. arabidopsidis* [86]. However new multigene phylogenies show that the *Peronospora* lineage has speciated after the divergence of *Phytophthora* clade 7 from clade 1 and 2. Notably, these three clades all contain species with duplicated genomes [2, 5, 6, 87]. This would suggest that an ancestral whole genome duplication before this speciation point would also apply to *Peronospora*, so that duplication cannot account for the difference in genome size. The availability of more *Peronospora* genomes for comparisons asks for a reevaluation of the timing of the duplication and subsequent speciation events.

Biologically, the question of how *Peronospora* species can be host-specific and obligate biotrophic while maintaining only a small and compact genome is interesting. It is argued that the trend in filamentous phytopathogens is towards large genomes with repetitive stretches to enhance genome plasticity [85]. Plasticity may enable host jumps and adaptations that favor the species for survival over species with small, less flexible genomes [85]. The reduced genomes of *Peronospora* species show an opposing trend that cannot be attributed to their obligate biotrophic lifestyle alone, as it is not evident in *Plasmopara* species (75 Mb – 9 2 Mb) [5, 88]. Sequencing of multiple races of the same *Peronospora* species may shed light on genome plasticity at the species level.

### Secretome reflects biotrophic lifestyle

#### Evolving biotrophy

The biotrophic lifestyle has emerged on several independent occasions in filamentous plant pathogens, in several branches of the tree of life. Convergent evolution is thought to be the main driving factor behind the development of biotrophy in such distantly related organisms [89]. However, it was shown that horizontal gene transfer can also occur between fungi and oomycetes, resulting in 21 fungal proteins in the secretome of *Hp. arabidopsidis*. Out of these 21 proteins, 13 were predicted to secreted, indicating that horizontal gene transfer may effect a species pathogenicity and interaction with the host [90, 91].

It was proposed that the critical step for adopting biotrophy in filamentous phytopathogens is the ability to create and maintain functional haustoria [92]. To do so, a species needs to be able to avoid host recognition or suppress the host defense response. A proposed mechanism for avoidance of host recognition is the loss of proteins involved in cell wall degradation, as evidenced by the reduction of cell wall degrading enzymes in mutualistic species compared to biotrophs [93]. In this and other studies, we find a reduction of the number of cell wall degrading enzymes in obligate biotrophic species compared to hemibiotrophic *Phytophthora* species (S3 Fig) [27]. This is true for all three obligate biotrophic groups in this study (*Hyaloperonospora/Peronospora, Plasmopara* and *Albugo*) although the difference is less clear in *Plasmopara*. Possibly this reduction is the result of a similar selection pressure to reduce recognition by the host plant in the biotrophic species, where the hemibiotrophic nature of the interaction between host and Phytophthora allows for slightly less caution in recognition avoidance.

The other mechanism of establishing a strong interaction is suppression or avoidance of the host defense response. Biotrophic infections are often accompanied by co-infection of species that are unable to infect the plant in the absence of the biotroph, indicating efficient defense suppression [92, 94]. We found enhanced numbers of secreted serine proteases (PF00082) (suppression) and reduced numbers of proteins with PAN/Apple domains that are known to be recognized by the plant immune system.

While the expansion of host translocated RxLR effectors is evident in both hemibiotrophic and biotrophic species, their numbers are smaller in secretomes of obligate biotrophs. Crinkler effectors are especially reduced in secretomes of biotrophic species. As opposed to RxLR effectors, CRNs are an ancient class of effectors that are known to induce cell death. Obligate biotrophic species presumably lost them as they are not beneficial for their survival.

In this study we first showed that the CAT tool performs well for taxonomic filtering of eukaryotic contigs. We provided a clean reference genome of the oldest known isolate of the spinach infecting downy mildew, *Pfs1.* In a comparative approach, we found that the secretomes of the obligate biotrophic oomycetes are more similar to each other than to more closely related hemibiotrophic species when comparing the presence and absence of functional domains, including the host translocated effectors. We conclude that adaptation to biotrophy is reflected in the secretome of oomycete species.

## Acknowledgments

We wish to thank Topsector Horticulture and Starting Materials (TKI) for funding the project. BED was supported by the Netherlands Organisation for Scientific Research (NWO) Vidi grant 864.14.004. We thank Ronnie de Jonge (Utrecht University) for useful input for the orthology analysis, Bjorn Wouterse for helping out with the comparative and statistical analysis, and the Utrecht Sequencing Facility for providing sequencing service and data. Utrecht Sequencing Facility is subsidized by the University Medical Center Utrecht, Hubrecht Institute, Utrecht University and The Netherlands X-omics Initiative (NWO).

## Supporting information

### Supplemental Figures

**S1 Fig. GC plot of various oomycete assemblies on contigs larger than 1kb.**

**S2 Fig. PCA on absolute numbers of secreted CAZyme domains.**

**S3 Fig. Secreted cell wall degrading proteins (CAZymes).** Numbers (a) of literature curated plant cell wall degrading enzymes per species. (b) The same data represented as fraction of the total number cell wall degrading protein domains per species.

**S4 Fig. PCA on absolute numbers of secreted Pfam domains.**

**S5 Fig. Secreted pathogenicity associated Pfam domains.** Occurrence of Pfam domains known to be involved in pathogenicity within the secretome of each species. Figure (a) shows the absolute number of Pfam domains, while (b) shows the number relative to the total number of Pfam domains per species.

### Supplemental Tables

**S1 Table. Species used for comparative secretomics.**

**S2 Table. Comparison of conserved eukaryotic genes for different oomycetes and the *Pfs1* assembly using BUSCO.**

**S3 Table. Repeat elements in the *Pfs1* genome.** Repeat elements identified in the *Pfs1* genome, for each repeat type the total numbers and percentage are shown. In addition, also a detailed annotation for each repeat element is provided.

**S4 Table. Genome sizes and repeat content of different assembled oomycete genomes.**

**S5 Table. Putative annotations of the *Pfs* proteins as obtained with ANNIE.** In addition, the presence of a N-terminal signal peptide for secretion, WY motif, TM motif and overlap with a repeat region are listed for each protein coding gene.

**S6 Table**. **Overview of the host translocated effectors (RxLR and CRN) identified in the genome of *Pfs1*.** Also, their respective functional domains and locations are listed per effector. Selected effectors that were used in the gene intergenic distance analysis are listed in the second tab.

**S7 Table. Secreted CAZyme domains per species.**

**S8 Table. Secreted Pfam domains per species.**

